# The KH domains of FMRP differentially regulate neuronal granule formation, dynamics, and function in *Drosophila*

**DOI:** 10.1101/2021.06.01.446613

**Authors:** Emily L. Starke, Scott A. Barbee

## Abstract

Fragile X Syndrome (FXS) is the most prevalent cause of inherited mental deficiency and is the most common monogenetic cause of autism spectral disorder (ASD). Here, we demonstrate that disease-causing missense mutations in the conserved K homology (KH) RNA binding domains (RBDs) of FMRP cause defects in its ability to form RNA transport granules in neurons. Using molecular, genetic, and imaging approaches in the *Drosophila* FXS model system, we show that the KH1 and KH2 domains of FMRP regulate distinct aspects of neuronal FMRP granule formation, dynamics, and transport. Furthermore, mutations in both KH domains disrupt translational repression in cells and the localization of known FMRP target mRNAs in neurons. These results suggest that the KH domains play an essential role in neuronal FMRP granule formation and function which may be linked to the molecular pathogenesis of FXS.

## INTRODUCTION

Fragile X Syndrome (FXS) is the most common cause of inherited intellectual disability in humans (Santoro et al., 2012). Typically, FXS is caused by epigenetic silencing of the *FMR1* gene due to a long CGG repeat expansion in the 5’UTR, resulting in hypermethylation of the *FMR1* locus and subsequent transcriptional silencing (Pieretti et al., 1991). This results in loss of expression of the encoded Fragile X Mental Retardation Protein (FMRP), an evolutionarily conserved RNA-binding protein (RBP) that binds to many mRNAs in the mammalian brain. FMRP is best characterized as a translational repressor (Richter and Zhao, 2021). In this role, FMRP associates with diverse ribonucleoprotein particles (RNPs) including RNA transport granules, P-bodies (PBs), and stress granules (SGs) (Lai et al., 2020). In neurons, FMRP-containing RNA transport granules (hereafter called “FMRP granules”) are actively transported in both axons and dendrites (Antar et al., 2005; Antar et al., 2006; Dictenberg et al., 2008; Li et al., 2009). These granules carry translationally repressed mRNAs to synapses where they are derepressed in response to synaptic activity. Local translation of critical mRNAs at synapses is essential for long-term synaptic plasticity and is defective in FXS.

The role of FMRP in translation and mRNA transport in the context of neurons remains enigmatic. In the soma, FMRP binds to translationally repressed target mRNAs and associated RBPs. These RNPs merge and are remodeled to allow for rapid, motor-dependent transport within neurites (El Fatimy et al., 2016). FMRP granules belong to a diverse class of membraneless organelles (MLOs) that form through liquid-liquid phase separation (LLSP) (Banani et al., 2017). This process is driven by weak, multivalent interactions between protein and RNA components (Mittag and Parker, 2018; Van Treeck and Parker, 2018). Weak interactions allow MLOs to be highly dynamic and to rapidly assemble and disassemble in response to local signals. In the case of FMRP, posttranslational modification of its C-terminal intrinsically disordered region (IDR) can reversibly control its phase separation *in vitro,* a process that correlates with translational repression (Tsang et al., 2019). This is an attractive model to explain how FMRP granules might assemble, deliver translationally repressed mRNAs to the synapse, and then regulate their local translation in response to activity. However, it is unclear whether the IDR acts alone or together with structured RBDs to regulate FMRP granules *in vivo*.

Although the most common cause of FXS is loss-of-function, advances in gene sequencing have led to the discovery of FXS-causing missense mutations in the *FMR1* gene (Suhl and Warren, 2015). Two mutations located in structured N-terminal RBDs of FMRP and have been functionally characterized (De Boulle et al., 1993; Feng et al., 1997; Myrick et al., 2015a; Myrick et al., 2015b; Prieto et al., 2021; Zang et al., 2009). The Gly266Glu (G266E) and Ile304Asn (I304N) mutations are in K-homology domains (KH1 and KH2 respectively) which bind to “kissing-complex” tertiary motifs or distinct sequence elements (GACR, ACUK, and WGGA) within target mRNAs (Ascano et al., 2012; Darnell et al., 2005; Ray et al., 2013). The latter are ubiquitous sequences in mammalian transcripts (Suhl et al., 2014). The analysis of these mutations has begun to uncover novel functions for *FMR1*. For example, both the G266E and I304N disrupt the ability of FMRP to bind to specific target mRNAs and to associate with polysomes suggesting that the KH domains are important for translational regulation (Myrick et al., 2014; Zang *et al*., 2009). However, the precise role these domains play in FMRP granule formation, dynamics, and function in the context of neurons remains unknown.

Studying FMRP function in mammals is genetically complicated due to the presence of two autosomal paralogs, FXR1P and FXR2P, which have some functional redundancy with FMRP (Agulhon et al., 1999; Wan et al., 2000; Zhang et al., 1995). In contrast, *Drosophila* has a single *dFmr1* gene and the dFMRP protein shares significant sequence identity with mammalian FMRP, particularly within the RBDs (Wan *et al*., 2000). *Drosophila* FMRP granules are also compositionally like those observed in mammalian neurons(Barbee et al., 2006; Cougot et al., 2008). Importantly, *Drosophila* has proven to be an excellent model system in which to study FXS because *dFmr1* mutants recapitulate many FXS phenotypes (Drozd et al., 2018). Here, we have introduced analogous mutations into the KH1 and KH2 domains of dFMRP (G269E and I307N respectively) and examined FMRP granules in *Drosophila* Schneider (S2) cells and primary neurons. Analysis of these mutants has revealed distinct differences in the requirement for KH1 and KH2 in the formation and dynamics of FMRP granules. Interestingly, both FXS-causing KH mutations result in FMRP granules that are significantly more dynamic. This is in opposition to mutations in other RBPs that drive the formation of solid-like aggregates associated with neurodegenerative disorders. Finally, the KH mutations differentially impact the function of FMRP in translational repression and RNA transport. Our findings provide new biological insight into the normal function of FMRP in cells and neurons and into the specific molecular and cellular processes that are dysregulated in FXS.

## RESULTS

### The KH domains are required to form FMRP granules in cells

The C-terminal IDR of mammalian FMRP is necessary and sufficient to drive the formation of phase-separated liquid droplets *in vitro* (Tsang *et al*., 2019). However, the dependency of the IDR in FMRP granule formation in cells has yet to be elucidated. To address this, we first developed dFMRP deletion and mutant transgenes (Figure 1A). Unless otherwise noted, EGFP was fused to the N-terminus to visualize granules. Overexpression of GFP-tagged wild-type dFMRP (WT-FMRP) in *Drosophila* larval motor neurons (MNs) replicated published results with untagged dFMRP indicating that GFP does not significantly interfere with protein function (Figure S1) (Zhang et al., 2001).

**Figure 1:**
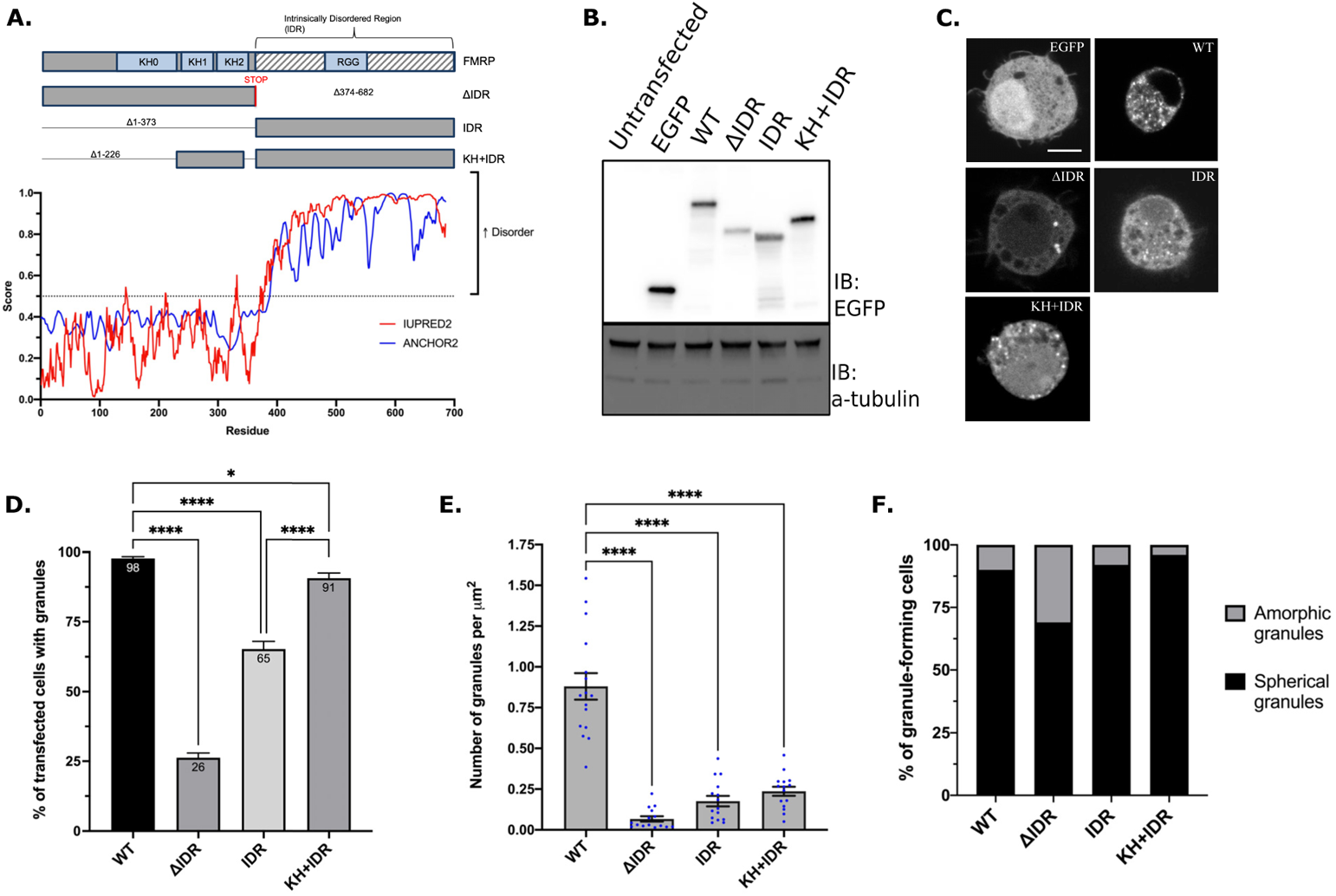
The KH domains and lOR interact to regulate FMRP granule formation. (A) Schematic of dFMRP showing each of the main RBDs in light blue boxes and lOR indicated by grey and white stripes (top). lOR mutants denote amino acid deletions with lines. Both KH1 and KH2 domains are fused to the lOR in the KH+IDR mutant. Disorder plot aligned with the wild-type dFMRP protein show that the C-terminus is entirely disordered as predicted by IUPRED2 and ANCHOR2 72. (B) Western blot analysis of EGFP (GFP) and a-tubulin (loading control) protein levels in transfected cells. (C) Representative images of GFP-FMRP mutant granule phenotypes in transfected S2R+ cells. Scale bar = 2µm. (D) Percentage of trans fee ted cells forming GFP-FMRP granules. Data are presented as mean ± S.E. of three independent experiments (approximately 100 cells per experiment; one-way ANOVA). (E) Quantification of the number of granules counted with in a cell, which was normalized to cell area in µm^2^ (mean ± SE; n=15 cells; Brown-Forsyth test). (F) Quantification of the two major morphological phenotypes observed in lOR mutants (n= 100 cells). In 0 and E, * p<0.05; **** p<0.0001.

The C-terminus of dFMRP is predicted to be disordered, indicating it may play an important role in promoting LLPS (Figure 1A). We first transfected *Drosophila* S2R+ cells with WT-dFMRP, dFMRP_ΔIDR_ (ΔIDR), and dFMRP_IDR_ (IDR) (Figure 1B-C). As shown previously, 98% of cells transfected with WT-FMRP form numerous small round granules (Figure 1C-D) (Gareau et al., 2013a; Gareau et al., 2013b). We found that the IDR alone was sufficient to induce FMRP granules in 65% of transfected cells (Figure 1C-D). These granules were morphologically like WT-FMRP albeit less numerous (Figure 1E-F). Interestingly, the structured N-terminal domain of dFMRP (ΔIDR) was also sufficient to induce granule formation in 26% of cells (Figure 1C-D). These granules were less abundant, and many had an amorphic (non-circular) morphology (Figure 1E-F). This suggests that, unlike human FMRP *in vitro*, the C-terminal IDR of dFMRP is not essential for FMRP granule formation in cells (Tsang *et al*., 2019).

We speculated that the N-terminal KH1 and KH2 domains may act cooperatively with the IDR to regulate FMRP granule formation. Fusing the KH domains to the IDR (KH+IDR) significantly increased the number of cells containing granules (Figure 1D).

These foci were morphologically indistinct from WT-FMRP granules although the number of KH+IDR granules per cell did not increase significantly (Figure 1C and 1E-F). These data indicate that the propensity to form granules is enhanced by addition of the KH domains to the IDR. However, our results also suggest that additional elements in the N-terminus are likely to be involved in the control of FMRP granule formation.

### FXS-causing mutations in the KH domains disrupt granule formation

We next wanted to explore the contribution of the KH domains in FMRP granule formation. The G266E and I304N missense mutations in KH1 and KH2 of hsFMRP are predicted to disrupt the proper folding of each RBD and to disrupt functions of FMRP including mRNA-binding, AMPA receptor trafficking, and polysome association (Darnell *et al*., 2005; Myrick *et al*., 2014; Valverde et al., 2007). To address this, we made analogous point mutations in the KH domains of dFMRP (G269E and I307N), hereafter referred to as KH1* and KH2* (Figure 2A). In contrast to WT-FMRP, overexpression of the GFP-tagged KH1* and KH2* transgenes in larval MNs did not inhibit axon terminal growth at the neuromuscular junction (NMJ) (Figure S1) (Zhang *et al*., 2001).

**Figure 2:**
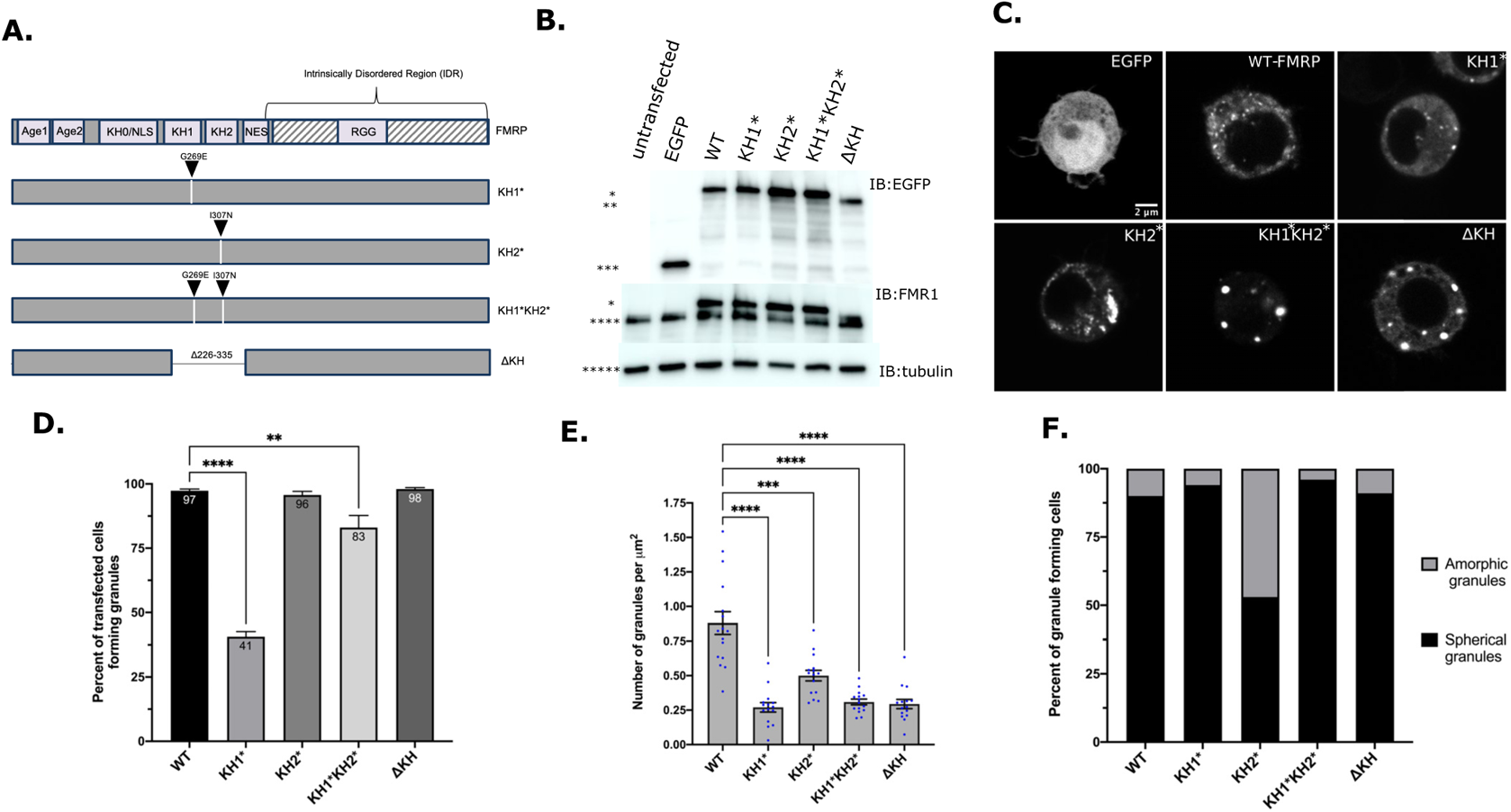
The KH domains differentially regulate FMRP granule formation. (A) Schematic representation of dFMRP variants used in th is study. Arrowheads indicate where analogous FXS-causing point mutations were made in dFMRP. Deletion of the KH1 and KH2 domains is annotated with a break in FMRP sequence. (B) Western blot analysis of EGFP (GFP), FMRP, and a-tubulin protein levels in transfected cells. α-tubul in was used as a loading control (*= 100 kDa, **= 90 kDa, ***= 30 kDa, ****= 80 kDa, *****= 60 kDa). (C) Representative images of cells transiently transfected with the indicated GFP-tagged FMRP constructs. Scale bar = 2 µm. (D) Percentage of transfected cells form ing GFP-FMRP granules. Data are presented as mean ± S.E. (approximately 100 cells per three experiments; one-way ANOVA). (E) Quantification of the number of granules per cell, which was normalized to cell area in µm^2^ (mean ± SE; n=15 cells each; Brown-Forsyth test). (F) Quantification of the two major morphological phenotypes observed (n=100 cells each). In D and E, **p<0.01, ***p<0.001. ****p<0.0001.

We transfected S2R+ cells with GFP-tagged constructs to determine the impact these mutations had on FMRP granule formation. Interestingly, we observed a > 2-fold decrease in the ability of cells expressing GFP-tagged KH1* to form granules relative to WT-FMRP, while KH2* had no effect (Figure 2C-D). This decrease cannot be explained by a difference in the expression levels of GFP-KH1* (Figure 2B). We also found that the number of granules per cell was significantly reduced by both mutations, although, granules were about twice as abundant in KH2* than KH1* (Figure 2E). Many KH2* granules also had an unusual, often large, amorphic structure while both WT-FMRP and KH1* granules were generally small and round (Figure 2C and 2F). These results suggest that both the KH1 and KH2 mutants alter normal FMRP granule formation. They also suggest that each KH domain control a different aspect of this process.

To examine the collective contribution of both domains, we made a G269E/I307N double mutant (KH1*KH2*) (Figure 2A). Most cells transfected with these constructs were able to form granules (Figure 2C-D). As with the individual KH mutants, transfected cells contained fewer granules per cell (Figure 2E). Interestingly, most KH1*KH2* granules that formed in cells were large and round indicating that they are different from those containing WT-FMRP, KH1*, or KH2* (Figure 2C and 2F). Morphologically these were like a published mutant where both KH domains have been deleted (ΔKH) (Gareau *et al*., 2013a; Gareau *et al*., 2013b). Together, these data suggest that the KH domains work together to restrict the size and shape of FMRP granules. However, we cannot rule out that the disruption of both KH domains is changing overall protein structure which is causing the formation of aggregates containing GFP-ΔKH and GFP-KH1*KH2* protein.

### FXS-causing mutations in the KH domains alter the dynamics of FMRP granules

A defining feature of phase separated RNPs in cells is the ability of components to rapidly shuttle between granules and the cytosol. The driving force underlying this process is multivalent interactions between protein and RNA components (Shin and Brangwynne, 2017). To examine whether the G269E and I307N mutations influenced dFMRP dynamics, we first conducted Fluorescence Recovery After Photobleaching (FRAP) experiments in S2R+ cells. The fluorescent signal of GFP-tagged WT-FMRP recovered to 82% with a t_1/2_ of 21.9s, in agreement with published results (Figure 3C-D) (Gareau *et al*., 2013b). We found that the exchangeable pool of KH1* and KH2* was like WT-FMRP (Figure 3C). However, KH1* and KH2* granules recovered more rapidly (t_1/2_ = 4.3 and 13.1s) suggesting these foci are much more dynamic than WT-FMRP (Figure 3A-B and 3D). Collectively, these data indicate that the individual KH mutations decrease the stability of FMRP in the mobile fraction. Finally, we examined the effect of removing or disrupting both KH domains. Compared to single mutants, KH1*KH2* significantly reduced dFMRP in the mobile fraction to 66% suggesting a large shift of FMRP into the non-dynamic immobile fraction (Figure 3C). Similar results were observed with the ΔKH mutant, however, recovery time of FMRP in the mobile fraction was significantly increased (t_1/2_ = 98.7s) (Figure 3D). These results further suggest that disruption of both the KH domains may lead to the strengthening of interactions or protein aggregation.

**Figure 3:**
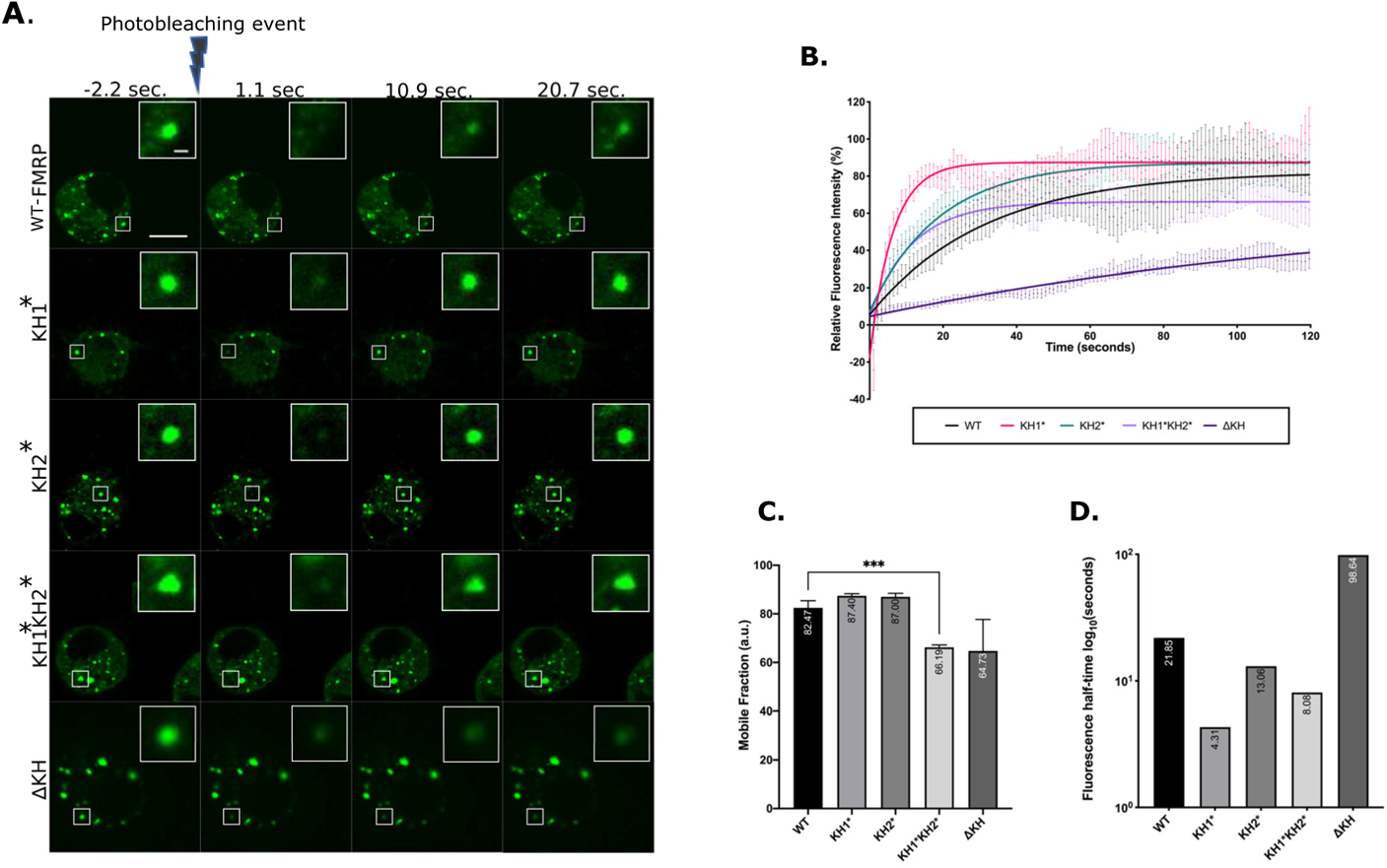
FXS-causing mutants alter FMRP granule dynamics in S2R+ cells. (A) Representa tive time-lapse FRAP images of FMRP-mutants pre- and post-bleaching. Scale bar in whole cell image = 5µm. Scale bar in zoomed-in granule image = 0.5µm. (B) Fluorescence recovery curves of FMRP-mutants over 120 seconds. Data points are mean ± SE. (C) Mobile fraction of FMRP mutant granules (mean ± SE; Brown-Forsyth test; ***p<0.001). a.u.=arbitrary units. (D) Quantification of the average time in 10910 (seconds), for granules to recover to half their final intensity (t_1/2_). For B, C, and D n= 17-21 granules.

### FXS-causing mutations alter the liquid-like properties of stress granules

In addition to neuronal granules (NGs), FMRP is a component of both SGs and PBs in neurons (Lai *et al*., 2020). Precisely how FMRP interacts with different populations of granules has yet to be elucidated. We hypothesized that mutations in the KH domains might disrupt the association of FMRP with these RNP populations. To study interactions with SGs, we co-transfected S2R+ cells with GFP-tagged FMRP constructs and mCherry-tagged Rasputin (Rin), the fly ortholog of G3BP1, a conserved marker for and modulator of SG assembly (Tourriere et al., 2003). In concordance with previous studies, overexpression of Rin induced SG formation in ~20% of unstressed transfected cells (Figure 4A and 4D) (Tourriere *et al*., 2003). Interestingly, we found that these Rin-positive granules always contained GFP-tagged FMRP and GFP-KH1* always colocalized with Rin (Figure 4A). They were also resistant to treatment with 1,6-hexanediol (1,6-HD), an aliphatic alcohol believed to interfere with weak protein-protein (π-π) and protein-RNA (π-cation) interactions required to form liquid-like MLOs (Kroschwald et al., 2017) (Figure 4D). As expected, arsenite-induced stress triggered the formation of cytoplasmic SGs (Figure 4B and 4E). Surprisingly, co-transfection with GFP-KH1* or GFP-KH2* significantly reduced the number of cells that formed Rin-positive SGs (Figure 4E; left). Moreover, the SGs that formed in cells co-transfected with KH mutants were more resistant to 1,6-HD treatment then with WT-FMRP (Figure 4E; right). Together, this suggests that the liquid-like nature of SGs is partially disrupted by the KH mutations.

**Figure 4:**
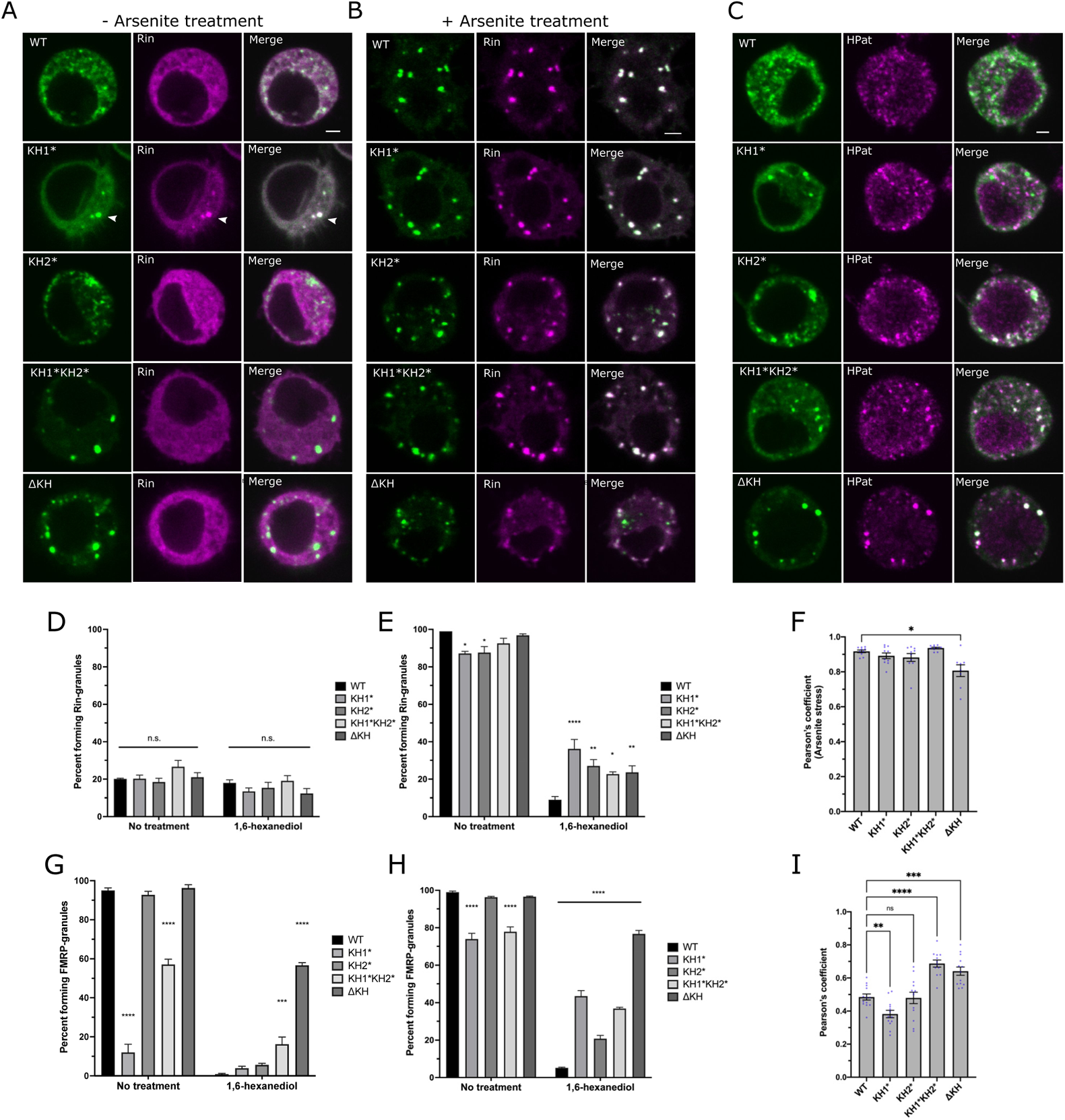
FXS-causing mutations alter SG dynamics and PB association. Representative images of S2R+ cells transfected with GFP-FMRP mutants (green) and Rin-mCherry (magenta) that are either not treated (A} or treated (B) with 0.5mM sodium arsenite for 45 minutes. Scale bars = 2µm. (C) Representative images of the localization of transiently transfected GFP-FMRP mutants immunostained against GFP (green) and HPat (magenta). Scale bar = 2µm. Percent of unstressed (D) or arsenite stressed (E) transfected cells forming Rin-positive SGs with or without 10% 1,6-HD treatment. Comparisons are made to WT-FMRP in each subcategory (mean ± SE; - 100 cells in triplicate; one-way ANOVA). (F) Average Pearson’s correlation coefficient between FMRP-mutants and the stress granule marker, Rin, in arsenite treated cells (mean ± SE of 8-10 cells; Brown-Forsyth test). Percentage of unstressed (G) or arsenite stressed (H) transfected cells forming FMRP granules with or without 10% 1,6-HD compared to WT­ FMRP (mean ± SE; - 100 cells in triplicate; one-way ANOVA). (I) Average Pearson’s correlation coefficient between FMRP-mutants and HPat (mean ± SE of 12-13 cells; one­ way ANOVA). In all graphs: * p<0.05, **p<0.01, ***p<0.001, ****p<0.0001.

Mammalian FMRP also accumulates in SGs under conditions of arsenite stress (Kim et al., 2006). In concordance with these results, all GFP-tagged dFMRP constructs colocalized strongly with Rin in stressed cells (Figure 4F). The number of cells containing KH mutant granules also increased in stressed cells but those with KH1* and KH1*KH2* were still significantly lower than WT-FMRP (Figure 4G-H; left). As with Rin, all KH mutant granules in stressed cells were significantly more resistant to 1,6-HD compared to WT-FMRP (Figure 4H; right). In the case of the KH1* and KH2* mutants, this was not likely due to the persistence of pre-existing FMRP granules because these nearly disappear in unstressed cells (Figure 4G; right). This further suggest that the liquid-like nature of SGs has been disrupted by the single KH mutants. The presence of KH1*KH2* and ΔKH granules in unstressed cells and their resistance to 1,6-HD treatment provides a third line of evidence suggesting that these mutations are causing FMRP to aggregate.

### The KH1 domain is required for the localization of FMRP to P-bodies

In addition to SGs, FMRP has been shown to colocalize with PB proteins in fly and mammalian neurons (Barbee *et al*., 2006; Cougot *et al*., 2008). Thus, we were next interested in determining if the KH mutants affected the ability of FMRP to interact with HPat/Pat1p, a highly conserved PB protein that colocalizes with FMRP in *Drosophila* neurons (Coller and Parker, 2005; Pradhan et al., 2012). To address this, we co-transfected S2R+ cells with GFP-tagged FMRP constructs and immunostained against HPat (Figure 4C). As expected, WT-FMRP overlapped moderately with HPat-positive granules (Figure 4I). In comparison, colocalization was significantly reduced in KH1* and most punctate GFP-KH1* failed to colocalize with punctate HPat (Figure 4C and 4I). Taken together, these data suggest that the KH1 domain is required for the association of FMRP with PBs. In contrast, KH1*KH2* and ΔKH caused the formation of larger granules that strongly colocalized with HPat, suggesting that PB proteins are recruited to these structures (Figure 4C and 4I). Based on these results and its propensity to form solid-like aggregates, we excluded ΔKH from subsequent analyses in neurons.

### The KH1 domain is required for FMRP granule formation in primary neurons

FMRP granules are important for the regulated trafficking of FMRP and specific RNA cargos in axons and dendrites (Antar et al., 2004; Antar *et al*., 2005; Antar *et al*., 2006; Davidovic et al., 2007; Dictenberg *et al*., 2008; El Fatimy *et al*., 2016). Based on our results in S2R+ cells, we asked if either of the KH domains played a role in the assembly or dynamics of FMRP granules in neurons. We generated inducible GFP-tagged dFMRP transgenic lines. First, we examined fly viability in a *dFmr1^Δ50M^/dFmr1^Δ113^* (*Δ50/Δ*113) loss-of-function genetic background when transgenes were expressed in larval motor neurons (*C380-Gal4, cha-Gal80* driver). As described, we found that the *Δ50/Δ*113 allele combination was viable (Morales et al., 2002). Surprisingly, motor neuron-specific expression of the GFP-KH1* and KH1*KH2* transgenes caused embryonic lethality in *Δ50/Δ*113 mutant flies (data not shown). As a result, we conducted all experiments below in a *Δ50/+* heterozygous background.

We next examined FMRP granules in 4-day old primary motor neuron cultures from dissociated larval ventral ganglia. All cells expressing WT-FMRP formed generally small, round granules in the soma (Figure 5A-B). In contrast, only 88% of cells expressing KH2* formed granules and, like what we observed in S2R+ cells, they were less numerous and sometimes formed amorphous structures (Figure 5A-B). Strikingly, GFP-KH1* and KH1*KH2* formed few granules in the soma of only 2% and 10% of neurons (respectively) suggesting that the KH1 domain is required to form FMRP granules *in vivo* (Figure 5A-B). The level of expression of each KH mutant was similar, indicating that this result was not likely due to reduced protein concentrations (Figure 5C). Both WT-FMRP and KH2* granules were also found in neurites (Figure 5D). However, there were significantly fewer GFP-KH2* granules outside of the soma (Figure 5D-E). Moreover, fewer mutant granules were found in distal regions of neurites (Figure 5F). Together, these data suggest that the KH2 domain is required for FMRP granule transport.

**Figure 5:**
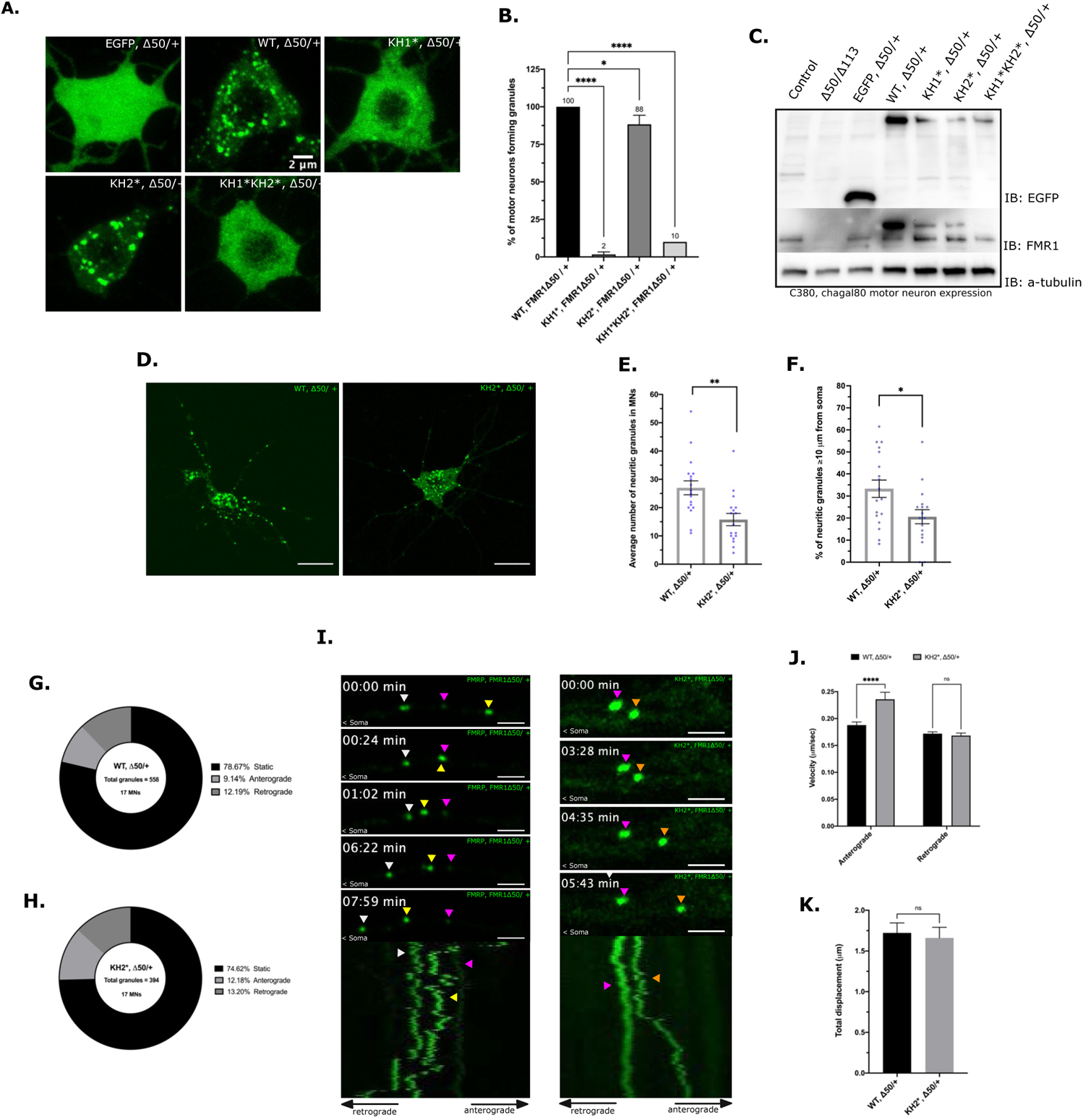
FXS-causing mutations disrupt NG formation and trafficking in neurons. (A) Representative images of major granule phenotype in primary neuron cell bodies. Expression of the indicated transgene was driven by *C380-Ga/4, cha - ga/80*. Scale bar = 2µm. (B) Percent of GFP-positive motor neurons forming FMRP granules in the *dFmr1 ^Δ^*^5 0/*+*^ background. Average is shown above respective bar (mean ± SE; 20 cells per triplicate, one-way ANOVA). (C) Western blot analysis of EGFP (top), FMRP (middle), and a,-tubulin (bottom) expression under the *C380-Ga/4, cha - ga/80* selective motor neuron driverin the larval CNS. (D) Representaitve imagesof *WT, Δ50/+* and *KH2*, L150/ +* pr ima ry MNs. Scale bar = 1Oµm. (E) Quantification of the average number of NGs within neurites of primary MNs (mean ± SE; 17-18 MNs, unpaired t test). (F) Percentage of neuritic granules that are distal (:2:10 µm) from the MN cell body (mean± SE; 17-18 MNs, unpaired t test). Pie charts representing the fraction of neuritic granules that remain stationary (static/oscillatory)or move in the anterograde or retrograde direction (relative to the cell body) in *WT, Δ50/ +* ( G) or *K H 2*, Δ50/ +* (H) primary neurons. Percentages are annotated in the legend for each chart (n= total granules in 17 MNs). (I) Time-lapse imagesand kymographs illustrating NG movements within neurites of WT (left panel) and KH2* (right panel) NGs. Images are oriented with the cell body on the left. Each granule is annotated with a colored arrowhead which corresponds with the traces in the kymograph. Scale bar= 2 µm. (J) Comparison of anterograde and retrograde velocities of motile WT-FMRP and KH2* NGs in neurites (mean± SE; 46-75 granules per category; two-way ANOVA). (K) Average total displacement (µm) of all motile WT and KH2* NGs (mean± SE; unpaired t test). In all graphs: * p<0.05, **p<0.01, ***p<0.001, ****p<0.0001.

### The KH2 domain is required for FMRP granule trafficking in neurites

We next asked if the KH mutations caused defects in the transport of FMRP granules. As GFP-KH1* and KH1*KH2* do not form any granules in neurites (Figure 5A), we focused on *KH2*,Δ50/+* compared to *WT-FMRP,Δ50/+.* Consistent with recent findings of GFP-tagged FMRP in hippocampal neuron dendrites, the majority of neuritic WT-FMRP granules were stationary (Figure 5G) (El Fatimy *et al*., 2016). Although, the number of mobile granules in KH2 mutants was not significantly different, slightly more KH2* granules were transported in the anterograde direction and the velocity of transport was significantly increased (Figure 5H-J). Despite this increase in net velocity, total granule displacement was not significantly altered (Figure 5K). These data, in conjunction with the reduction in the number of FMRP granules found in distal neurites (Figure 5D-F), suggest that the KH2 mutant granules have significant transport defects.

### The KH2 domain regulates the dynamics of FMRP in neuronal granules

We next performed FRAP analysis in primary *Drosophila* motor neurons, looking at somatic and neuritic granules as two separate populations (Figure 6A-D). In agreement with published results, the mobile fraction of WT-FMRP was significantly lower in both the soma and neurites (33% and 35% respectively) of cultured neurons than in S2R+ cells (82%) suggesting that a larger proportion of WT-FMRP in NGs is found within the non-dynamic, immobile fraction (Figure 6E and 3C) (Estes et al., 2008; Gareau *et al*., 2013b). While the size of the immobile fraction of NGs in both compartments was similar, the recovery time of WT-FMRP in the mobile pool was about 2-fold slower in neuritic granules (t_1/2_ = 75s) than in those found in the soma (t_1/2_ = 37s) (Figure 6E-F). Additionally, both had slower recovery times than what we saw in S2R+ cells (Figure 3C-D). Together, these data suggest that wild-type FMRP granules in neurons are less dynamic than those that form in S2R+ cells. There were two observations with GFP-KH2* granules that support the conclusion that this mutation alters FMRP granule dynamics in neurons. First, the amount of KH2* found in the mobile fraction was significantly smaller in both somatic and neuritic NGs (Figure 6E). Second, the recovery time of KH2* in the mobile fraction was reduced (t_1/2_ = 5s) (Figure 6F). These data further indicate that the composition and dynamics of FMRP granules have been significantly altered by the KH2 mutation and suggest that the KH domains are required to stabilize FMRP granules in neurons. Based on these data, we propose that the disruption of this stability is contributing directly to the transport defects seen in KH2* granules (Figure 5D-K).

**Figure 6:**
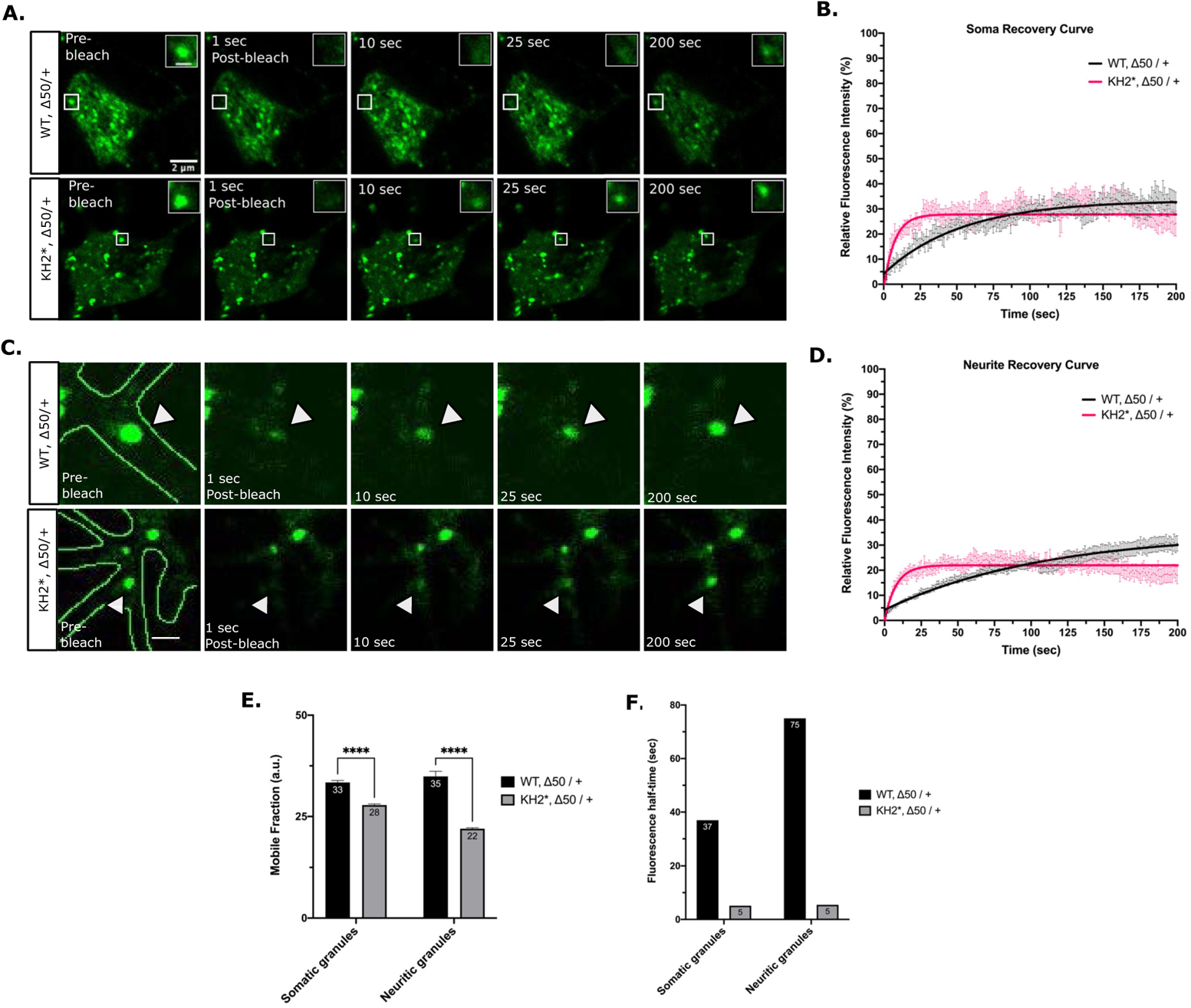
FXS-causing mutations in FMRP disrupt NG dynamics in neurons. (A) Representative FRAP time lapse images of somatic NGs pre- and post-bleaching event. Scale bar in whole cell images = 2 µm. Scale bar in zoomed-in granule images = 1 µm. (B) Fluorescence recovery curves of somatic NGs over 200 seconds (mean ± SE; n 9 gra nules). (C) Representative FRAP time-lapse images of neuritic NGs pre- and post-bleaching event. Neu rites are outlined in green in the pre-bleach image, arrow heads point to the bleached granule. Scale bar = 1µm. (D) Fluorescence recovery curves of neuritic NGs showing fluorescence intensity relative to the initial pre-bleach intensity over 200 seconds (mean ± SE; n 13 granules). (E) Quantification of the a verage mobile fractions of somatic (left) or neuritic (right) mobile fraction of WT and KH2* NGs. a.u.=arbitrary units; p < 0.0001. (F) Quantification of the fluorescence half-time (t_1/2_) of so m at ic or neu r itic WT an d K H 2 * NGs in seco n ds. For E and F n 9 granules.

### The KH domains are essential for the translational repression activity of FMRP

We next wanted to examine the role of the KH domains in regulating the translational repression activity of dFMRP. Both KH1* and KH2* have been shown to disrupt the association of mammalian FMRP with polysomes suggesting that the KH domains are important for translational repression (Darnell *et al*., 2005; Feng *et al*., 1997; Myrick *et al*., 2014; Zang *et al*., 2009). Moreover, the KH domains of dFMRP bind directly to the 80S ribosome and can block elongation *in vitro* (Chen et al., 2014). To further examine the role of the KH domains in translational control, we used a λN-based tethering assay in S2 cells, where the 3’UTRs of known dFMRP target mRNAs were fused to the 3’ end of a firefly luciferase reporter, or FLuc (Figure 7A). To eliminate RNA binding as a mechanism, we first tethered λN-tagged FMRP constructs directly to the reporter via a 5X tandem BoxB sequence that was inserted into the heterologous *SV40* 3’UTR. The WT-FMRP and KH2* constructs were both able to repress translation (Figure 7B). In contrast, the ability of KH1* to repress reporter expression was significantly disrupted, suggesting that the KH1 domain is required for repression activity.

**Figure 7:**
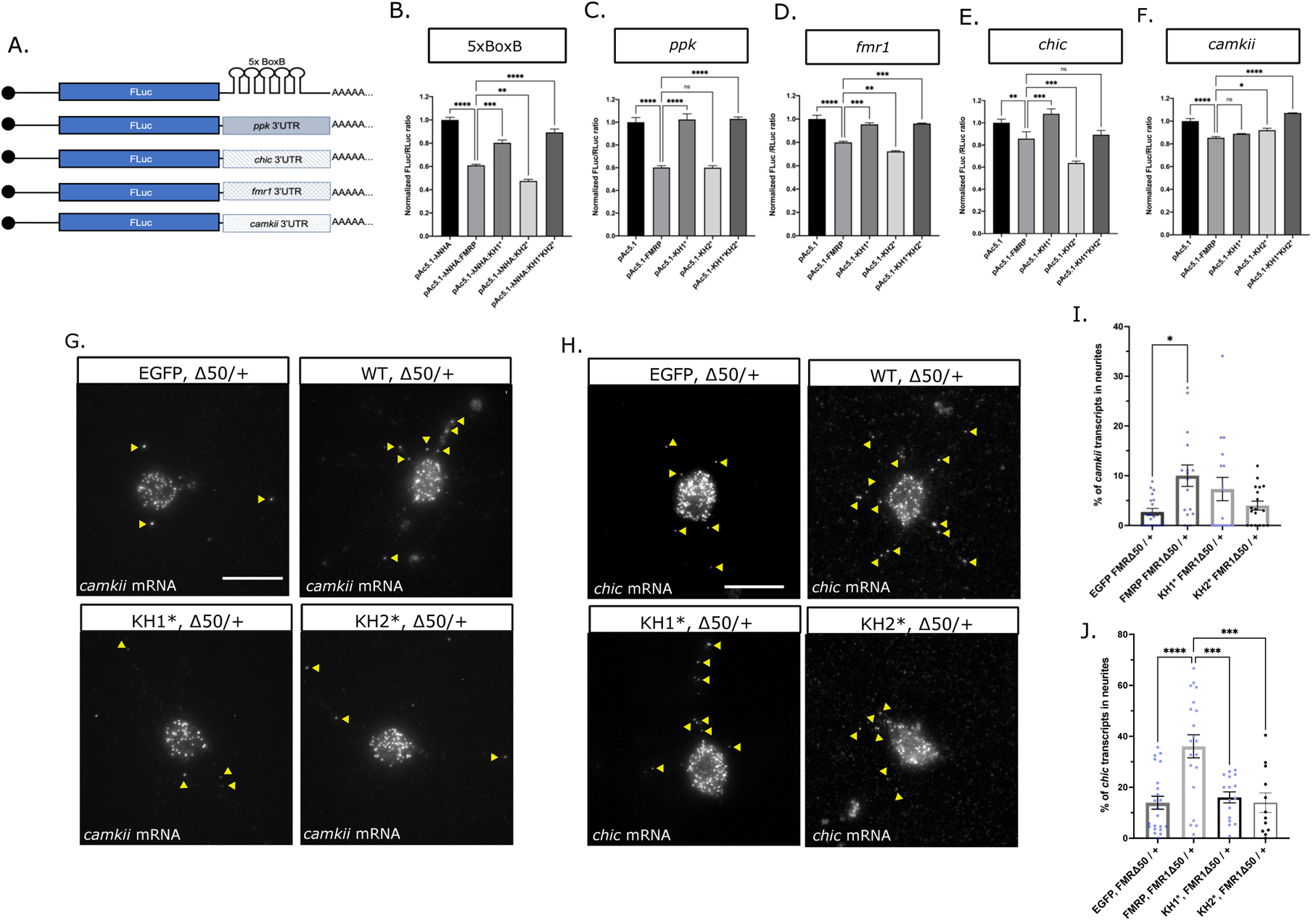
FXS-causing mutations disrupt translation and RNA transport. (A) Diagram of the FLuc reporters used in this study fused to the *SV40 3’UTR* containing the 5xBoxB sequence or to the 3’UTR’s of known mRNA targets of dFMRP. luciferase assays of (B) λN:HA-tethe r ed FMRP-mutantsrepression of the 5xBoxB FLuc reporter or the untethered FMRP-mutants repression of Flue fused to the (C) *pickpocket (ppk)*, (D) *fmr1*, (E) *chickadee (chic)* or (F) *camkii* 3’UTR. FLuc/RLuc ratios were normalized to empty vector ratios. Graph shows repression of the Flue reporter by empty vector or FXS-causing point mutants compared to *pAc5.1-λNHA:FMRP(B) or pAc5.1-FMRP (C-F)* (mean ± SE; one-way ANOVA). Representative images of *camkii* (G) or *chic* (H) mRNA smFISH in primary MNs. Yellow arrowheads in images are distinguishing transcripts found in neurites. Scale bars = 10μm. Quantification of the average number of *camkii* (I) or *chic* (J) transcripts in neurites of each of the FMRP mutants (mean ± SE of 11-25 MNs; unpaired t test). In all graphs: * p<0.05, **p<0.01, ***p<0.001, ****p<0.0001.

Next, we determined whether untethered dFMRP could repress the translation of Fluc by binding to the 3’UTRs of known targets mRNAs and if either of the KH domains were required for this to occur. We replaced the *SV40* 3’UTR containing the BoxB repeats with the 3’UTRs from mRNAs encoding for: 1) the degenerin/epithelial sodium channel (DEG/ENaC) family member, *pickpocket* (*ppk*); 2) the Ca^2+^/calmodulin-dependent protein kinase II, *camkii*; 3) the profilin ortholog, *chickadee (chic)*; and 4) its own mRNA, *fmr1* (Reeve et al., 2005; Sudhakaran et al., 2014; Xu et al., 2004; Zhang *et al*., 2001) (Figure 7A). As with the BoxB reporter, the KH1 domain was required to regulate repression of the *ppk*, *chic,* and *fmr1* reporters as repression was ameliorated in KH1* and KH1*KH2* (Figure 7B-E). The efficiency of repression of each of these reporters by WT-FMRP and KH2* was variable, likely due to differences in the ability of these proteins to interact with or bind to the 3’UTR. In contrast, the *camkii* reporter was different in that repression is slightly but significantly derepressed by KH2* and KH1*KH2* indicating that the KH2 domain is required for repression (Figure 7F). Collectively, these data indicate that FXS-causing mutations in the KH1 and KH2 domains can differentially regulate translational repression. The derepression of translation in KH1* correlates with defects in its ability to form FMRP granules in cells and neurons (Figure 2 and 5).

### The KH domains are required for the transport of FMRP target RNAs

NGs are specialized MLOs within neurons that serve to transport translationally silent mRNAs between the soma and axonal or dendritic compartments (Formicola et al., 2019). Therefore, we next asked if *KH1*,Δ50/+* or *KH2*,Δ50/+* neurons had defects in mRNA localization. To address this, we used single-molecule fluorescence *in situ* hybridization (smFISH) to quantify transcripts in the soma and neurites. We focused on two mRNA targets of dFMRP in flies, *chic* and *camkii*, both of which have been shown to interact with dFMRP-containing NGs in primary motor neurons (Barbee *et al*., 2006; Estes *et al*., 2008). Their translational repression is also regulated by the KH1 and KH2 domains (Figure 7E-F). We find that there are significantly more *camkii* and *chic* transcripts in *WT-FMRP,Δ50/+* neurites compared to controls suggesting that dFMRP promotes the transport of both mRNAs to neurites (Figure 7G-J). The percentage of *chic* transcripts found in neurites is significantly reduced in both KH1* and KH2*. In contrast, localization of *camkii* is not affected by either mutation. Taken together, these data suggest that the *chic* and *camkii* mRNAs are differentially localized to neurites through a KH domain-dependent and independent mechanism (respectively).

## DISCUSSION

FMRP has been implicated in a growing number of biological processes in neurons ranging from the control of translation and RNA transport to the regulation of RNA editing, splicing, genome stability, and ion channel function (Davis and Broadie, 2017). Because the vast majority of FXS cases are caused by loss of FMRP expression, it has been difficult to determine which of these functions are involved in FXS pathophysiology. In contrast, the study of disease-causing missense mutations in FMRP has allowed for the isolation of specific protein functions that may be contributing to FXS phenotypes (Suhl and Warren, 2015). Here, we provide multiple lines of evidence that FXS-causing mutations in the KH domains differentially affect FMRP granule assembly, dynamics, and function in *Drosophila* neurons. First, an FXS-causing missense mutation in the KH1 domain disrupts the ability of FMRP to form granules in primary neuron cell culture (Figure 5A-B). In contrast, KH2 mutants form granules in the soma but their transport to distal neurites is reduced (Figure 5D-F). Second, we find that KH2* significantly decreases the amount and stability of FMRP found in the mobile fraction of neuronal FMRP granules (Figure 6E-F). Third, the KH domains are differentially required to regulate the translation of reporters for known target mRNAs (Figure 7B-F). Finally, both KH domains promote the localization of specific target mRNAs to neurites (Figure 7G-J). Translational repression and RNA transport are processes that have been directly attributed to FMRP-containing RNA granules in neurons (Richter and Zhao, 2021).

The disordered C-terminus of mammalian FMRP is necessary and sufficient to drive the formation of phase-separated droplets *in vitro* (Tsang *et al*., 2019). Furthermore, phosphorylation patterns of amino acids in this region control the propensity to phase separate in the presence of RNA and to regulate rates of deadenylation and translation within these condensates (Kim et al., 2019). Collectively, these data suggest that regulation of FMRP phase separation might be a simple mechanism to allow for the delivery of translationally repressed mRNAs to synapses and to control their local translation in response to activity. Through our analysis of *Drosophila* FMRP granules in cells, we show that the C-terminal IDR is sufficient to regulate granule formation (Figure 1D). However, granule formation is significantly enhanced by the addition of the structured KH domains (Figure 1D). This is consistent with current models suggesting that MLO formation *in vivo* is driven by multivalent interactions between protein and RNA components (Mittag and Parker, 2018; Van Treeck and Parker, 2018). Increased valency provides a scaffold of *cis*-acting binding sites that allow for interactions with multiple *trans*-acting partners, allowing interacting molecules to compartmentalize within the cell (Banani et al., 2016; Li et al., 2012). Published findings also show that different types of NGs are generally more stable than other types of MLOs (Chae et al., 2010; Cougot *et al*., 2008; Formicola *et al*., 2019; Gopal et al., 2017; Vijayakumar et al., 2019). In support of this, we find that WT-FMRP granules are significantly less dynamic in primary neurons than in S2R+ cells (Figure 3 and 7). This stability is likely to be necessary so that FMRP granules can resist the shear stress associated with active transport.

While the precise RNA binding sites are a source of debate, the KH domains clearly bind with different specificity and/or affinity to target mRNAs (Athar and Joseph, 2020). Therefore, the simplest explanation for the role of the KH domains in regulating FMRP granule formation and dynamics is that they are increasing valency through interactions with RNA. Why does the KH1 mutation have a more significant impact on FMRP granules? It is possible that the KH1 domain is contributing disproportionately strong (or numerous) interactions with mRNAs which is shifting the critical concentration needed for granule formation. Interactions occurring via KH1 could be shifting the concentration threshold required to promote FMRP granule formation. For example, multivalent interactions between RNA and proteins such as hnRNPA1 and FUS, influence LLPS by shifting the phase boundary and requiring lower protein concentrations to initiate demixing (Molliex et al., 2015; Schwartz et al., 2013; Tsang *et al*., 2019).

Our analysis of KH domain function also reveals a link between FMRP granule formation and translational repression in neurons. A significant fraction of mammalian and *Drosophila* FMRP interacts with polysomes and this association is disrupted by the G266E and I304N mutations (Feng *et al*., 1997; Ishizuka et al., 2002; Myrick *et al*., 2014; Zang *et al*., 2009). One mechanism by which FMRP blocks translation is by interacting with stalled polysomes (Darnell et al., 2011). More specifically, the KH domains of dFMRP interact directly with the peptidyl site of ribosomes (Chen *et al*., 2014). This suggests a mechanism by which FMRP stalls translational elongation by sterically inhibiting tRNA entry. The non-disease associated KH1 mutant (I244N) and KH2* (I307N) both disrupt binding affinity, with the I244N mutation having a stronger effect (Chen *et al*., 2014). Interestingly, analysis of FMRP granules in mouse brain homogenates by electron microscopy found that FMRP and ribosomes both localize to a subset of neuronal RNA transport granules (El Fatimy *et al*., 2016). These data led the authors to propose that FMRP granules form from stalled polysomes. A prediction from this model is that disruption of FMRP-ribosome association would negatively impact the formation of a subset of FMRP granules. Our data provide support for this model as we show that KH1* and KH2* both disrupt FMRP granules, with the KH1 mutant having a stronger effect (Figure 2C-D and 5A-B). We also demonstrate that the KH1 domain is required to repress the translation of most translational reporters tested, including when FMRP was directly tethered to a heterologous 3’UTR (Figure 7B-E). Additional studies would be required to determine if there is a direct connection between polysomes, ribosome stalling, and FMRP granule formation in *Drosophila* neurons. This would also help to clarify whether FMRP granules are a cause or consequence of translational repression.

In cultured mouse neurons, the specificity of FMRP for mRNAs targeted for transport to neurites is conferred by interactions between its C-terminal RGG domain and G-quadraplex sequences within the 3’UTRs of localized mRNAs (Goering et al., 2020). This study also found that the FXS-causing KH2 mutant (I304N) which disrupts ribosome association retained the ability to promote the localization of G-quadraplex-containing transcripts to neurites. In contrast, we show that both the KH1 and KH2 mutants fail to promote the localization of *chic* (but not *camkii*) to neurites in cultured *Drosophila* neurons (Figure 7G-J). The long *chic* 3’UTR (but not *camkii*) contains G-rich sequences predicted to fold into quadraplexes (data not shown) suggesting that localization of these fly mRNAs may be regulated by a G-quadraplex-independent mechanism. This might be due to our limited sample size as mouse studies focused on a global analysis of mRNA localization (Goering *et al*., 2020). Alternatively, the control of mRNA transport by *Drosophila* FMRP may be mechanistically different. The RGG domain found in dFMRP is not highly conserved and it fails to bind with high affinity to the *sc1* RNA, a G-quadraplex-containing target for human FMRP (Darnell et al., 2009). Unlike in mouse studies, we also observe both transcript- and KH domain-specific coupling of translational repression and mRNA localization in *Drosophila* neurons (Figure 7) (Darnell *et al*., 2009). It is possible that this represents a *bona fide* difference between fly and mammalian FMRP.

The role of FMRP granules in the transport and translation of target mRNAs in neurons is complex because their organization, dynamics, and function are regulated by multivalent interactions involving both structured and unstructured protein domains. Our data support a model where FMRP granules are metastable, solid-like, MLOs. This is a state that is likely necessary to allow for their active transport in neurites without the loss of RNA and protein components. We demonstrate that the KH1 and KH2 domains are required to maintain FMRP granule integrity in both S2R+ cells and neurons. FXS-causing mutations disrupt this equilibrium, leading to the formation of MLOs that are unstable. This is distinctly different from disease-causing mutations in IDR-containing RBPs that have been linked to neurodegenerative disease such as TDP-43, FUS, or hnRNP-A1, which lead to the formation of metastable, then stable, pathological inclusions (Molliex *et al*., 2015; Patel et al., 2015). That said, our studies have revealed novel granule phenotypes associated with FXS-causing KH mutations. First, both KH mutations disrupt the sensitivity of SGs to 1,6-HD, suggesting that their biophysical properties have been altered (Figure 4D-H). KH1* has a more significant impact on this phenotype than KH2*. Second, MN-specific genetic “rescue” experiments with KH1* causes lethality in an otherwise viable *dFmr1* loss-of-function background suggesting that G266E may be an uncharacterized gain-of-function mutation (data not shown). Finally, the KH1 mutation also nearly abolishes FMRP granule formation in neurons. These results may explain why the G226E mutation causes such a severe form of FXS (Myrick *et al*., 2014).

## METHODS

### Fly stocks and husbandry

In all experiments, both male and female flies were used. All lines were incubated at 25°C with 12-hour light/dark cycles and 60% humidity on standard Bloomington medium. Fly lines used and made in this study are listed in Table S1. Plasmid constructs for the generation of transgenic lines were constructed as described below. All transgenes were injected into fly strain 24485 for *PhiC31-*mediated integration into the same site on chromosome III. All transgenes were recombined with the *dFmr1^Δ50M^* null allele.

### Cell culture

S2 and S2R+ cells were maintained at 24°C with ambient humidity in a dark incubator in M3+BPYE media. S2-DRSC cells were used for luciferase reporter assay experiments. S2R+ cells were used for imaging experiments due to their higher propensity to adhere to and flatten out on poly-D-lysine coated cover slips. Cells and media recipes were obtained from the *Drosophila* Genomics Resource Center (DGRC). Primary motor neurons were cultured from wandering 3^rd^ instar larvae using a protocol adapted from Barbee et al. 2006. For each genotype, ventral ganglia from ten third instar larvae were dissected, dissociated, and then grown in complete M3+BPYE media (M3+BPYE supplemented with 50 μg/mL insulin) for 3 to 5 days prior to imaging.

### Cloning and site directed mutagenesis

The backbone vector for all S2 cell culture plasmids is a modification of pAc5.1 (ThermoFisher Scientific, Waltham, MA) which drives expression from the fly actin 5C gene. For imaging of FMRP, the coding region of the *FMR1-RD* cDNA (LD09557; DGRC) was PCR amplified and cloned downstream of EGFP in the pAc5.1B-EGFP vector (a gift from Elisa Izaurralde; Addgene plasmid #21181) using the *HindIII* and *EcoRI* restriction sites to make pAcB5.1-EGFP-FMRP. The pAc5.1:EGFP-FMRP-IDR, pAc5.1:EGFP-FMRP-ΔIDR, pAc5.1:EGFP-FMRP-KH:IDR and pAc5.1:EGFP-FMRP-ΔKH plasmids were all constructed by PCR amplification of the indicated sequence followed by cloning into pAc5.1B-EGFP. Primer sequences and restriction sites are listed and described in Table S2. The *Drosophila* G269E and I307N mutations were generated using the Q5 Site-Directed Mutagenesis Kit (New England Biolabs, Ipswitch, MA). Mutagenesis primers were designed using the “substitution” feature in NEBaseChanger v1.2.9 (New England Biolabs). Mutagenesis occurs at nucleotide 805-807 (GGA→ GAA) and nucleotide 868-870 (ATC→ AAC) in the KH1 and KH2 domain, respectively. Each mutation was introduced directly into the pAc5.1B:EGFP-FMRP vector. The KH1*KH2* double mutant was created by introducing both mutations sequentially (KH1* followed by KH2*).

Plasmids for the generation of transgenic fly lines were all constructed as follows. The EGFP-FMRP sequence for WT-FMRP and each mutant construct were PCR amplified from their respective pAc5.1 vector using primer sequences listed and described in Table S2. Products were then cloned directionally into the 5’-*KpnI* and 3’-*XbaI* sites of pUAST-attB. Transgenic flies were generated as described above.

To construct the pAc5.1-Rin-mCherry plasmid, we first generated a pAc5.1B-MCS-(Gly_4_Ser)_3_-mCherry plasmid. The Rin cDNA was obtained by reverse-transcriptase PCR from RNA extracted from wild-type (*CantonS*) adult flies. Total RNA was extracted using the Direct-zol RNA kit (Zymo Research, Irivne, CA) and cDNA was generated using the RNA to cDNA EcoDry™ Premix Oligo dT kit (Takara Bio USA, Moutain View, CA). Rin was PCR amplified using primers listed and described in Table S2 and cloned into the *HindIII* and *EcoRI* restriction sites of pAc5.1-MCS-(Gly_4_Ser)_3_-mCherry to generate the final pAc5.1:Rin-(Gly_4_Ser)_3_-mCherry vector.

To construct plasmids for the tethered FLuc reporter assays, the wild-type dFMR1 and KH1*, KH2*, or KH1*KH2* mutants were PCR amplified from their respective pAc5.1B-EGFP vector and cloned into the *HindIII* and *EcoRI* sites of the pAc5.1-lambdaN-HA vector (a gift from Elisa Izaurralde; Addgene plasmid #21302) (Behm-Ansmant et al., 2006). For the untethered reporter assays, the indicated dFMR1 constructs were cloned into the MCS of pAc5.1B lacking the lambdaN-HA tag. The BoxB reporter was pAc5.1C-Fluc-Stop-5BoxB (a gift from Elisa Izaurralde; Addgene plasmid #21301) (Behm-Ansmant *et al*., 2006). To construct the *camkii* and *Fmr1* reporters, the *SV40* 3’UTR and the 5xBoxB sequence was removed by digesting pAc5.1C-Fluc-Stop-5BoxB with *EcoRI* and *XhoI*. The long 3’UTR isoform of *camkii* and *Fmr1* were amplified by RT-PCR (as described for Rin) using primers listed and described in Table S2. The 3’UTRs were then cloned into the cut plasmid via Gibson Assembly using the Gibson Assembly Master Mix (New England Biolabs). The 3’UTRs of *ppk* and *chic* were amplified by RT-PCR, cloned into pENTR D-TOPO, and into the pAc5.1-FLuc2 [dPolyA] RfA destination vector as we have previously described (Nesler et al., 2013).

### S2 cell transfections

DNA transfections were performed with the Effectene Transfection Reagent kit (Qiagen). Transient transfections were performed following the manufacturer’s protocols. For fluorescence experiments, DNA constructs were transfected into S2R+ cells in 12-well plates. Cells were transfected with 0.5 μg of each construct except for pAc5.1B-EGFP:FMRP:KH1* in which 0.75 μg of DNA was used. Transfected cells were grown for ~ 72 hours before imaging. For luciferase assays, S2-DRSC cells in 24-well plates were transfected with 0.025 µg of the FLuc reporter plasmid, 0.1 µg of the *Renilla* luciferase (RLuc) transfection control plasmid, and 0.25 µg of either the pAc5.1/pAc5.1-λN control vector or the pAc5.1:FMRP/pAc5.1-λN:FMRP wild-type or mutant constructs. Transfected cells were grown for ~ 72 hours prior to analysis in luciferase experiments.

### S2 cell imaging and granule analysis

S2R+ cells were transferred to poly-D lysine coated #1.5 cover glass bottom dishes (Cellvis, Mountain View, CA) within two hours of imaging. In all experiments, images were obtained using an Olympus FV3000 scanning confocal microscope with a 100x (NA 1.4) objective digitally zoomed to 2.95x (the optimal setting per the Fluoview software). To quantify the number of granules formed by each construct, 15 cells were analyzed in a single experiment. The single z-plane where the nucleus took up the largest cell area was imaged. Punctate areas of fluorescence intensity above background were considered to be granules and were quantified using the Cell Counter plugin in ImageJ2/FIJI. For each image, cell diameter (through the longest axis in the same z-plane) and granule number were collected. From this, the number of granules per cell area (μm^2^) was recorded. To count the number of transfected cells able to form granules, ~ 100 cells per FMRP construct were manually identified at the microscope. The total number of cells that formed granules out of all EGFP-expressing cells were quantified. Cells were scanned in z to make sure that granules in all planes were identified. The number of cells that formed granules was divided by total number of transfected cells for three separate experiments. To compare the propensity of the different mutants to form circular granules, ~ 100 granule forming transfected cells were identified at the microscope. The number of cells forming circular and amorphic (non-circular) granules were counted. In most cases, cells that formed amorphous granules also contained circular granules, however, cells that formed any number of amorphous granules were categorized as amorphic.

### Fluorescence Recovery After Photobleaching (FRAP) experiments

For all FRAP experiments, 17-21 EGFP positive granules were imaged using an Olympus FV3000 scanning confocal microscope with a 100x (NA 1.4) objective digitally zoomed to 2.95x (optimal settings). Granules selected for FRAP analysis generally fit into the “spherical” category. Granules selected for analysis in primary neurons were clearly in the cell body or neuritic compartments. In all cases, granules were photobleached with the lowest laser intensity necessary to completely bleach the granule, ranging from 2.44-10% 488nm laser power for 500-1,000 milliseconds. Two pre-FRAP images were collected and images were captured every 1.0878 seconds pre and post bleaching for a total of 200 frames (~ 218 s). For image analysis, images were initially processed in ImageJ2/FIJI) and analyzed essentially as described (Cheney et al., 2017). A ROI was manually traced in each FRAP movie for 1) the bleached granule, 2) an unbleached granule and 3) diffuse cytoplasmic staining for background. As necessary, the ROI was moved if/when granules moved in x/y out of the initially set ROI. If a granule clearly dropped out of the focal plane, it was excluded from analysis. The mean fluorescent intensity was obtained for each frame. The following was calculated: 1) Photobleach Correction Value (PCV), in which the average intensity of the pre-bleach unbleached granule was divided by the value for each subsequent unbleached granule, 2) Corrected Average Intensity (CAI), where the bleached granules mean intensity was multiplied by the PCV, 3) Background Corrected Fluorescence Intensity (BCFI), where the CAI was subtracted by the average intensity of the background ROI, and 4) the Final Corrected Value (FCV) which was calculated by dividing each BCFI by the initial BCFI value and multiplying by 100 to get a normalized fluorescence intensity profile. The FCVs from each movie were then analyzed a one-phase association, nonlinear fit in Graphpad Prism to calculate the fluorescence recovery curve, mobile fraction, and half-life.

### Immunocytochemistry

S2R+ cells were transferred to poly-D lysine coated #1.0 cover glass bottom dishes (Cellvis) and allowed to settle for ~ 20 minutes. Cells were then fixed with 4% PFA for 10 minutes followed by a 5-minute incubation in ice cold methanol. Cells were washed three times for 5 minutes in PBS, permeabilized in PBST for 10 minutes, and then blocked in PBST with 2% BSA and 5% normal goat serum for 30 minutes. Cells were then incubated with primary antibodies overnight at 4°C, washed three times for 5 minutes in PBS, and then incubated for 1 hour in secondary antibodies at room temperature. Finally, cells were washed with PBS and then mounted in DAPI-Fluoromount-G Clear Mounting Media (ThermoFisher Scientific). EGFP:FMRP constructs were visualized by GFP fluorescence. For HPat, rabbit anti-HPat (1:1,500) (Pradhan *et al*., 2012) primary and goat anti-rabbit Alexa-567 (1:500) secondary antibodies were used.

### Arsenite and 1,6-hexanediol treatment

For SG experiments, cells were co-transfected with the indicated pAc5.1B:EGFP-FMRP construct and pAc5.1-Rin-mCherry. About 72 h post transfection, cells were treated with 0.5mM sodium arsenite in M3+BPYE media for 45 minutes to induce SG formation. Non-stressed cells went through the same procedure but were incubated in media that did not contain sodium arsenite. For colocalization analysis, cells were immediately fixed in 4% PFA for 10 minutes, incubated with ice cold methanol for 5 minutes and then washed 3 times for 5 minutes in PBS. Preparations were then mounted on coverslips as described above. FMRP and Rin were visualized by imaging GFP and mCherry fluorescence respectively. For the analysis of granules with treated with 1,6-HD, cells were co-transfected with pAc5.1:EGFP-FMRP constructs and pAc5.1-Rin-mCherry. Cells were subjected to arsenite stress (or not stressed) as described above. Following the 45-minute incubation (+/- sodium arsenite), cells were treated with M3+BPYE media (+/- sodium arsenite) containing 10% 1,6-HD. Approximately 100 live transfected cells were analyzed for the presence or absence of FMRP or Rin granules, in triplicate.

### Colocalization analysis

To determine the degree to which FMRP mutants colocalized with SGs or PBs, 12-13 images were analyzed in ImageJ2/FIJI using the JACoP plugin (Bolte and Cordelieres, 2006). Images were cropped to the smallest area possible to eliminate colocalization outside of the cell of interest and images for FMRP/HPat were background subtracted to a rolling ball radius of 50 pixels to account for the higher degree of HPat background staining. In JACoP, Pearson’s coefficient results were recorded for analysis.

### Western blotting

For the detection of EGFP-FMRP construct levels in S2R+ cells, transfected cells were harvested at three days post-transection from a 6-well plate. Cells were scraped and resuspended by pipetting up and down and 1.5 mL of cells were spun down at 1,000x G for 5 minutes at 4°C. Cells were then resuspended in 400 μL of 2x Laemmli sample buffer + β-mercaptoethanol on ice. For westerns conducted on *C380, cha-Gal80/+; UAS:EGFP-FMRP, FMR1^Δ50M^/+* larvae ectopically expressing the wild-type and FXS-causing point mutants, 5 ventral ganglia (including optic lobes) were dissected and added to 100μL of 2x Laemmli sample buffer + β-mercaptoethanol on ice. Then, the ventral ganglia were homogenized in a 1.6 mL microcentrifuge tube for 30 seconds on ice using a hand-held homogenizer. Homogenates were incubated on ice for 3 minutes, and then processed as indicated above for S2R+ cells. For both assays, the primary antibodies used were mouse anti-dFMR1 (1:3,000; 6A15; Abcam, Cambridge, UK), rabbit anti-EGFP (1:2,000; Proteintech Group, Rosemont, IL), and mouse anti-ɑ-tubulin (1:1,000; 12G10; Developmental Studies Hybridoma Bank, Iowa City, IA). Secondary antibodies used were horse anti-mouse HRP or goat anti-rabbit HRP which were both diluted 1:1,000 in block (Cell Signaling Technology, Danvers, MA).

### Primary motor neuron imaging and neurite transport analysis

For analysis of *C380, cha-Gal80/+; UAS:EGFP-FMRP, FMR1^Δ50M^/+* larvae expressing either wild-type FMRP or the KH2 mutant, primary larval motor neurons were collected and cultured as described above. At 4 days post-harvest, live neurons were imaged using an Olympus FV3000 scanning confocal microscope with a 100x (NA 1.4) objective. For soma imaging, images were digitally zoomed to 2.95x for optimal resolution and a z-stack was obtained with 0.39μm slices through the entire soma. Images were presented as Z-projections made using ImageJ2/FIJI. For neurite transport analysis, live cells were imaged with a 100x objective (NA 1.4) digitally zoomed to 1.79x so branching neurites could be imaged. Movies were collected containing four 0.39μm z-slices, over 100 frames (8:04 minutes). Movies were then analyzed using the Kymolyzer plugin in ImageJ2/FIJI from which granule velocities and directionality were obtained, using a lower speed limit set to the pixel size (0.138μm) (Basu et al., 2020). To calculate the average number of neuritic granules in primary motor neurons, the max-intensity Z-projection of the first time point was used (Frame 1). The number of granules in the soma and neurites were manually counted using the Cell Counter plugin for ImageJ2/FIJI. Additionally, the proportion of neuritic granules 10μm or further from the cell body was determined from these images. The scale was globally set, and a symmetrical circle was drawn tightly around each cell using the oval selection tool in ImageJ, containing as much of the cell as possible. The diameter of each cell body was recorded in μm. The center of each circle was determined and marked using the Pencil tool. A line was drawn to the center of each granule within neurites and distance from the cell body was recorded in μm. These were subtracted by the radius for their respective cell body to obtain the final distance.

### Luciferase reporter assays

Luciferase reporter assays were done essentially as we have described (Nesler *et al*., 2013). At three days post transfection, 75 µL of resuspended cells were added in three technical replicates to a 96-well white, flat bottom polystyrene assay plate (Costar, Washington, DC). FLuc and RLuc expression was measured using the Dual-Glo Luciferase Assay System kit (Promega, Madison, WI) per the manufacturer’s instructions and luminescence detected using a Synergy™ HTX Multi-Mode Microplate Reader (BioTek, Winooski, VT). FLuc values were normalized to corresponding RLuc values. The FLuc/RLuc ratio for each experiment was normalized to its respective control. The experiment was repeated three times for each FLuc reporter.

### Single molecule FISH and FISH-quant image analysis

For analysis of *C380, cha-Gal80/+; UAS:EGFP-FMRP, FMR1^Δ50M^/+* larvae expressing either wild-type FMRP or the KH2 mutant, primary larval motor neurons were collected and cultured as described above. Custom Stellaris® FISH Probes were designed against *Drosophila melanogaster camkii* and *chic* by utilizing the Stellaris^®^ FISH Probe Designer (Table S3; Biosearch Technologies, Hoddesdon, UK). Primary motor neurons were hybridized with the indicated Stellaris FISH Probe set labeled with either Quasar-570 or 670 (Biosearch Technologies) following the manufacturer’s instructions. Briefly, at 3-4 days post culturing, cells growing on #1.5 cover glass were washed in PBS. Cells were then incubated in fixation buffer (3.7% formaldehyde in PBS) for 10 minutes at room temperature, then washed twice in PBS. To permeabilize cells were immersed in 70% ethanol at 4°C for at least 1 hour and up to a week. Ethanol was aspirated and cells were washed in Stellaris Wash Buffer A for 5 minutes, then hybridized with the indicated probe(s) in a dark, humid hybridization chamber at 37°C for 5-16 hours. Probes were used at a final molarity of 0.125μM in Stellaris Hybridization buffer. Hybridization buffer was aspirated, and cells were incubated with Wash Buffer A twice at 37°C for 30 minutes, then washed with Stellaris Wash Buffer B for 5 minutes at room temperature. Buffer was aspirated and Vectashield Mounting Medium was added to the #1.5 cover glass in the imaging dish and a clean coverslip was placed on top and sealed with clear nail polish. Imaging dishes were stored in the dark at −20°C for up to 2 days before imaging on an ONI Nanoimager S. Approximately 15 cells were imaged per genotype using the widefield microscopy application. To detect smFISH probes, cells were exposed to 7% 570- or 640-laser power for 1,500 milliseconds. Z-projection was obtained with 0.2 μm slices through the entire cell. EGFP-FMRP was imaged sequentially which allowed us to distinguish the soma and neurites from background. To analyze smFISH images, we used the FISH-Quant Matlab application to detect, localize and quantify mRNA in primary motor neurons (Mueller et al., 2013). Motor neuron soma and neurites were outlined individually by hand, which allowed us to differentiate mRNAs residing within the soma and neurites.

### Quantification and statistical analysis

All data were initially recorded in Excel (Microsoft) and then graphed and analyzed in Prism version 9.0.2 (GraphPad). Results were considered statistically significant if p<0.05. Error bars throughout the study indicate mean ± SEM. n.s. = not significant, * p<0.05, ** p<0.01, *** p<0.001, and **** p<0.0001. Outliers were identified and removed using ROUT method in Prism. Statistical tests and sample sizes for each experiment are indicated within the corresponding figure legend and/or in methods section.

## AUTHOR CONTRIBUTIONS

Designed experiments: ELS, SAB. Performed experiments: ELS Analyzed the data: ELS, SAB. Wrote the paper: ELS, SAB

## ACKNOWLEDGMENTS

We thank members of the S.A.B. lab for useful discussions. S2 cell lines and cDNAs were obtained from the *Drosophila* Genomics Resource Center which is supported by NIH grant 2P40OD010949. Fly lines were obtained from the Bloomington *Drosophila* Stock Center which is funded by NIH grant P40OD018537. We thank Samantha Patterson for assistance with FRAP experiments, Sarala Pradhan for help with developing and testing the FMRP translational reporter assay, and Navneeta Kaul for assistance constructing several of the *pAc5.1-FLuc 3’UTR* reporters. We are also grateful to the Knoebel Center for Healthy Aging for use of the ONI NanoImager for smFISH experiments. We acknowledge funding for this study to S.A.B. from NIH grant R15MH114019 and from the Assistant Secretary of Defense for Health Affairs PRMRP grant W81XWH2110026.

