## Supplemental Data for "The KH domains of FMRP differentially regulate neuronal granule formation, dynamics, and function in *Drosophila*"

### SUPPLEMENTAL METHODS

#### Larval NMJ immunohistochemistry and morphological analysis

Third instar larval body wall preps for NMJ analysis were dissected in ice cold calcium-free Jan and Jan buffer (130 mM NaCl, 5 mM KCl, 36 mM sucrose, 5 mM HEPES [pH 7.3], 4 mM MgCl2, and 0.5 mM EGTA) within 30 minutes on sylgard plates. Dissection buffer was removed and preps were fixed in 4% PFA for 20 minutes, then washed 3x 5 minutes in 1X PBS (pH 7.4) and permeabilized in 1X PBS (pH 7.4) + 0.1% Triton X-100 for 10 minutes. Preps were blocked in 1X PBS with 2% BSA and 5% normal goat serum with shaking for 30 minutes. Block was removed and primary antibody (mouse anti-DLG, 1:200) diluted in block was incubated overnight at 4°C. Preps were washed 6x 5 minutes in PBS and then incubated for 1 hour with secondary antibodies diluted in block at room temperature (goat anti-mouse Alexa 488 or 567 at 1:500 & Alexa 649-conjugated anti-HRP at 1:500). Preps were washed in 1X PBS and then mounted on slides in DAPI-Fluoromount-G Clear Mounting Media. All imaging was done on an Olympus FV3000 scanning confocal microscope using 20X and 60X objectives to image the NMJ at muscles 6/7 in abdominal section 3 (N.A. 0.85 and 1.42, respectively). When shown, maximum Z projections were assembled from 0.4µm optical sections. All post-hoc image processing was done using Fiji in ImageJ2. For morphological analysis of larval NMJs, between 10-17 images were examined per experiment, in which 1s, 1b and axon terminals were manually counted at muscles 6 and 7 (m6/7) in abdominal segment 3 (A3) using the Cell Counter plugin in Fiji. To account for muscle area differences between genotypes which effects NMJ size, synaptic bouton numbers were normalized to muscle surface area (MSA). MSA was calculated by outlining both m6/7 using the freehand selection tool in ImageJ2/Fiji and recording the calculated muscle area. 1s and 1b bouton numbers were divided by the corresponding muscle area. These data were then normalized to the *C380-Gal4/+;; UAS-EGFP/+* controls. Data were collected and calculations were conducted in Excel and statistical analyses were performed in Prism.

Figure S1. KH domain mutants disrupt FMRP function when overexpressed in larval MNs

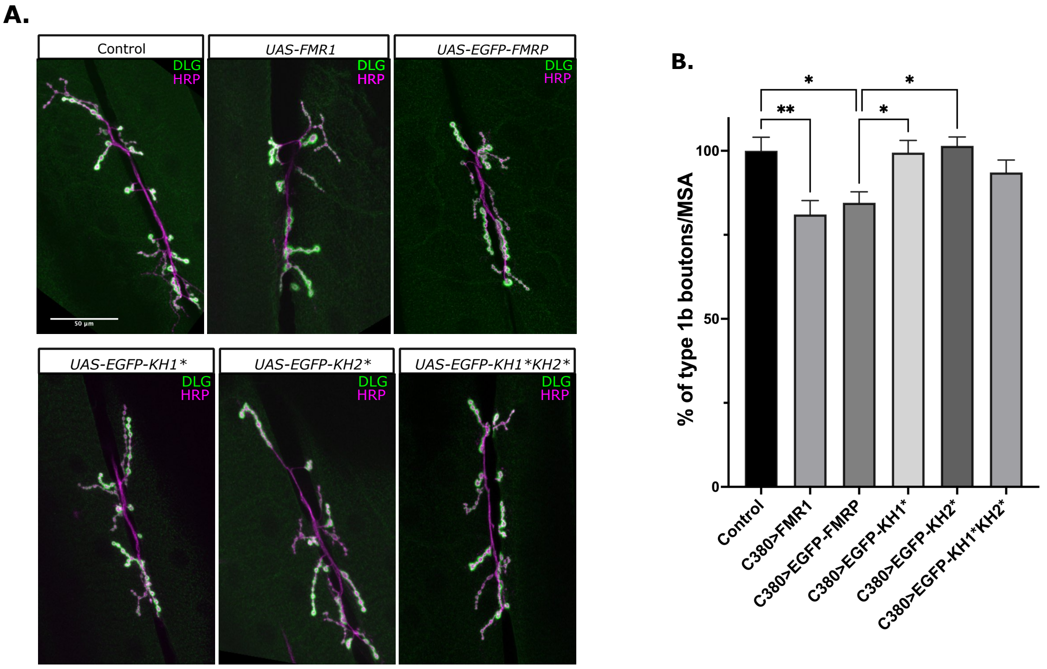

**Supplemental Figure 1: KH domain mutants disrupt FMRP function when overexpressed in larval MNs**
(A) Wandering third instar larval NMJs from *C380-Gal4* (Control), *UAS-FMR1*, and *UAS-EGFP:FMRP* mutants were stained with antibodies targeting the postsynaptic density marker, DLG (green) and the neuronal membrane marker, HRP (magenta). Maximum Z-projections of NMJs in abdominal segment 3 innervating body wall muscles 6/7 were analyzed. Scale bar = 50μm. (B) Percentage of type 1b bouton number normalized to the area of muscles 6/7 (in $\mu m^2$) was counted manually and compared with the control and EGFP:FMRP (mean ± SE; n=10-20 NMJs; one-way ANOVA). * $p<0.05$, ** $p<0.01$.

**Supplemental Table 1 – Fly lines**

| REAGENT or RESOURCE | SOURCE | IDENTIFIER |
|---|---|---|
| *D. melanogaster: BL Canton S* | Bloomington Drosophila Stock Center | BDSC:64349 |
| *D. melanogaster: C380-Gal4* | Bloomington Drosophila Stock Center | BDSC:80580 |
| *D. melanogaster: w$^{1118}$; UAS-FMR1* | Bloomington Drosophila Stock Center | BDSC:6931 |
| *D. melanogaster: w$^{1118}$;; FMR1Δ50M/TM6B,Tb$^{+}$* | Bloomington Drosophila Stock Center | BDSC:6928 |
| *D. melanogaster: w*;; FMR1$^{\Delta 113M}$/TM6B,T$^{b+}$* | Bloomington Drosophila Stock Center | BDSC:67403 |
| *D. melanogaster: C380-Gal4, cha-Gal80* | (Hartwig et al., 2008) | |
| *D. melanogaster: w$^{1118}$; wg$^{Sp-1}$/CyO, P{w$^{+}$mC=2xTb$^{1}$-RFP} CyO; MKRS/TM6B, Tb$^{1}$* | Bloomington Drosophila Stock Center | BDSC: 76359 |
| *D. melanogaster: C380-Gal4;; Sb/ TM6B,Ser* | This paper | |
| *D. melanogaster: pUAST-attB-EGFP* | This paper | |
| *D. melanogaster: pUAST-attB-EGFP:FMRP* | This paper | |
| *D. melanogaster: pUAST-attB-EGFP:FMRP:KH1** | This paper | |
| *D. melanogaster: pUAST-attB-EGFP:FMRP:KH2** | This paper | |
| *D. melanogaster: pUAST-attB-EGFP:FMRP:KH1*KH2** | This paper | |
| *D. melanogaster: pUAST-attB-EGFP:ΔKH* | This paper | |
| *D. melanogaster: w+; FMR1$^{\Delta 50M}$, pUAST-attB-EGFP/TM6BTb* | This paper | |
| *D. melanogaster: w$^{+}$; FMR1$^{\Delta 50M}$, pUAST-attB-EGFP:FMRP/TM6BTb* | This paper | |
| *D. melanogaster: w$^{+}$; FMR1$^{\Delta 50M}$, pUAST-attB-EGFP:FMRP:KH1*/TM6BTb* | This paper | |
| *D. melanogaster: w$^{+}$; FMR1$^{\Delta 50M}$, pUAST-attB-EGFP:FMRP:KH2*/TM6BTb* | This paper | |
| *D. melanogaster: w$^{+}$; FMR1$^{\Delta 50M}$, pUAST-attB-EGFP:FMRP:KH1*KH2*/TM6BTb* | This paper | |
| *D. melanogaster: w$^{+}$; FMR1$^{\Delta 50M}$, pUAST-attB-EGFP:ΔKH/TM6BTb* | This paper | |
| *D. melanogaster: C380,cha-Gal80;; TM6BTb/TM3BSb* | This paper | |
| *D. melanogaster: C380,cha-Gal80;; TM6BTb/FMR1$^{\Delta 113M}$* | This paper | |
| *D. melanogaster: C380-Gal4;; FMR1$^{\Delta 113M}$/TM6BTb* | This paper | |

**Supplemental Table 2 – Oligonucleotides**

| REAGENT or RESOURCE | SOURCE | IDENTIFIER |
|---|---|---|
| Oligonucleotides | | |
| PCR forward primer for genotyping fmr1 deletion: 5'-AAGGAAAAAAGCGGCCGCAAAGATATCGCGAAAATCCCCCCAG-3' | (Zhang et al., 2001) | |
| PCR reverse primer for genotyping fmr1 deletion 5'-CGGGATCCGTTATGCTACGTGAATAAATC-3' | (Zhang et al., 2001) | |
| Forward primer for amplifying the N-terminus of DmFMRP with a 5'-HindIII site: 5'-ACAAGCCAAGCTTTATGGAAGAT-3' | This paper | |
| Reverse primer for amplifying the C-terminal half of DmFMRP with a 3' EcoRI site: 5'-TCTGCAGAATTCTTAGGACGTG-3' | This paper | |
| Forward primer for amplifying the N-terminus of EGFP with a 5' KpnI site for cloning EGFP and all EGFP:FMRP mutants into the pUAST vector: 5'-GGTACCAACATGGTGAGCAA-3' | This paper | |
| Reverse primer for amplifying the C-terminus of EGFP with a 3' XbaI site for cloning EGFP into the pUAST vector: 5'-GTTCATCTAGACTACTTGTACAGCTCGTCCATGC-3' | This paper | |
| Reverse primer for amplifying the C-terminal portion of FMRP without the IDR to construct ΔIDR with a 3' EcoRI site: 5'-TACGGAATTCTTACTTCTCCTGACGCAACTGTT-3' | This paper | |
| Forward primer for amplifying the FMRP IDR with a 5' HindIII site: 5'-GTCAAAGCTTCGAGATTGATCAGCAGCTTC-3' | This paper | |
| Forward primer for amplifying the C-terminal half of DmFMRP to construct the KH-domain deletion(ΔKH) with a 5' BamHI site: 5'-ATGACGGATCCCTGGCGCATGTACCCTTTGT -3' | This paper | |
| Reverse primer for amplifying the N-terminal half of DmFMRP to construct the KH-domain deletion (ΔKH) with a 3' BamHI site: 5'-ATGACGGATCCCTCAACGTAGTTTCCACGGC-3' | This paper | |
| Forward primer for SDM of the KH1 domain in dmFMRP [Gly269Glu]: 5' CAAAATCAGCGAAGAGACCGAGG -3' | This paper | |
| Reverse primer for SDM of the KH1 domain in dmFMRP [Gly269Glu]: 5'-AATGTGCAGGACTTCTCC-3' | This paper | |
| Forward primer for SDM of the KH2 domain in dmFMRP [Ile307Asn]: 5'-GGGCGCATTAACCAGGAGATTG-3' | This paper | |
| Reverse primer for SDM of the KH2 domain in dmFMRP [Ile307Asn]: 5'-ATTCTTGCCAATCACCTTGC-3' | This paper | |
| mCherry amplification forward primer with 5' HindIII site): 5'-AGTACAAGCTTATGGTGAGCAAGGGCGAGGAG-3' | This paper | |
| mCherry amplification reverse primer with 3' BamHI site: 5'-AGTACGGATCCTTACTTGTACAGCTCGTCCATGCCG -3' | This paper | |

| | |
|---|---|
| Top primer for cloning (Gly4Ser)3 linker upstream of mcherry, containing a 5' ApaI site and 3' HindIII: 5'-CGGTGGAGGAGGCTCTGGTGGAGGCGGTAGCGGAGGCGGAGGGTCGA-3' | This paper |
| Bottom primer for cloning (Gly4Ser)3 linker upstream of mcherry, containing ApaI and HindIII sites: 5'-AGCTTCGACCCTCCGCCTCCGCTACCGCCTCCACCAGAGCCTCCTCCACCGGGCC-3' | This paper |
| Rasputin RT-PCR primer with 5' KpnI site: 5'-TGACATGGTCATGGATGCGACCCA-3' | This paper |
| Rasputin RT-PCR primer with in-frame stop codon and 3'-EcoRI site: 5'-ATACGAATTCGCGACGTCCGTAGTTGCCA-3' | This paper |
| CaMKII 3'UTR Gibson assembly primer for cloning into FLuc backbone vector cut with EcoRI and XhoI: 5'-CGGAAAGTCCAAATTGTAATGGGCATTAATCAATGGAATATAAAC-3' | This paper |
| CaMKII 3'UTR Gibson assembly primer for cloning into FLuc backbone vector cut with EcoRI and XhoI: 5'-CTTACCTTCGAATGGGTGACAAAATTGCATTATGCTTTGAATTC-3' | This paper |
| Forward restriction primer for cloning FMR1's 3'UTR containing the 5' EcoRI site: 5'-TACTGAATTCAGGAGCAACAGCTCACAG-3' | This paper |
| Reverse restriction primer for cloning FMR1's 3'UTR containing the 3' XhoI site: 5'-ATACCTCGAGGCTTGATGGTTTGTGTTTTG-3' | This paper |
| Forward primer for amplifying the *ppk* 3’UTR: 5’-CACCTCGATGGTCTTAAAGGCCGAAAG-3’ | This paper |
| Reverse primer for amplifying the *ppk* 3’UTR: 5’-GCGAACACATTTTTTATTGTCGTG-3’ | This paper |
| Forward primer for amplifying the *chic* 3’UTR: 5’-CACCCCGCTTCCGTGGTAGAGAAACT-3’ | This paper |
| Reverse primer for amplifying the *chic* 3’UTR: 5’-TGACTTTGGGAACCGCGATA-3’ | This paper |

**Supplemental Table 3 – smFISH probes**

| REAGENT or RESOURCE | SOURCE | IDENTIFIER |
|---|---|---|
| Oligonucleotides | | |
| camkii_1: 5’- GAAAAACGCGTACAGGCTGC-3’ Quasar 670 | LGC Biosearch Technologies | SS631302-01 |
| camkii_2: 5’- CCAACTCTTCTTTGATGTCG-3’ Quasar 670 | LGC Biosearch Technologies | SS631302-02 |
| camkii_3: 5’- GCAGCAAATTCAAAGCCAGT-3’ Quasar 670 | LGC Biosearch Technologies | SS631302-03 |
| camkii_4: 5’- CACTATGTTGGGATGGTGTA-3’ Quasar 670 | LGC Biosearch Technologies | SS631302-04 |
| camkii_5: 5’- CTCCTGTATACTGTCATGTA-3’ Quasar 670 | LGC Biosearch Technologies | SS631302-05 |
| camkii_6: 5’- AATGTGATGCATCAGCTTCT-3’ Quasar 670 | LGC Biosearch Technologies | SS631302-06 |
| camkii_7: 5’- CATTTTGGTGGCAGTGATTG-3’ Quasar 670 | LGC Biosearch Technologies | SS631302-07 |
| camkii_8: 5’- ATTCTCTGGTTTCAGATCTC-3’ Quasar 670 | LGC Biosearch Technologies | SS631302-08 |
| camkii_9: 5’- AGACCAAAGTCAGCGAGTTT-3’ Quasar 670 | LGC Biosearch Technologies | SS631302-09 |
| camkii_10: 5’- CTGATGATCGCCTTGAACTT-3’ Quasar 670 | LGC Biosearch Technologies | SS631302-10 |
| camkii_11: 5’- CTCCTTTTTCAATACCTCAG-3’ Quasar 670 | LGC Biosearch Technologies | SS631302-11 |
| camkii_12: 5’- AAGAATAACTCCACATGCCC-3’ Quasar 670 | LGC Biosearch Technologies | SS631302-12 |
| camkii_13: 5’- TGCTGATCTTCATCCCAAAA-3’ Quasar 670 | LGC Biosearch Technologies | SS631302-13 |
| camkii_14: 5’- ACGGATAATCATAAGCTCCC-3’ Quasar 670 | LGC Biosearch Technologies | SS631302-14 |
| camkii_15: 5’- TTTAGCTTCTGGAGTAACCG-3’ Quasar 670 | LGC Biosearch Technologies | SS631302-15 |
| camkii_16: 5’- GATGTTTTAAAGCCTCAGCT-3’ Quasar 670 | LGC Biosearch Technologies | SS631302-16 |
| camkii_17: 5’- CACACGTTCGCGTTGACAAA-3’ Quasar 670 | LGC Biosearch Technologies | SS631302-17 |
| camkii_18: 5’- CTTGAGACAGTCTACGGTTT-3’ Quasar 670 | LGC Biosearch Technologies | SS631302-18 |
| camkii_19: 5’- CGCCAACATTGTCGTAAGTA-3’ Quasar 670 | LGC Biosearch Technologies | SS631302-19 |
| camkii_20: 5’- GTTATCATACTTCTGCTCGA-3’ Quasar 670 | LGC Biosearch Technologies | SS631302-20 |
| camkii_21: 5’- GTTGATTCTTTGACCTGTGA-3’ Quasar 670 | LGC Biosearch Technologies | SS631302-21 |
| camkii_22: 5’- CGTCTTCAAGAGTAGTGCTA-3’ Quasar 670 | LGC Biosearch Technologies | SS631302-22 |
| camkii_23: 5’- GCCACTGTTAATTGCTTCAA-3’ Quasar 670 | LGC Biosearch Technologies | SS631302-23 |
| camkii_24: 5’- CAAAGGCAGTTAGATGCGGA-3’ Quasar 670 | LGC Biosearch Technologies | SS631302-24 |
| camkii_25: 5’- ATTCCTTCTACAAGGTTACC-3’ Quasar 670 | LGC Biosearch Technologies | SS631302-25 |

| | | |
|---|---|---|
| camkii_26: 5’- GCTTTGCAGTTTTTACCAAG-3’ Quasar 670 | LGC Biosearch Technologies | SS631302-26 |
| camkii_27: 5’- CTTCACCAAGTAAGTGCACA-3’ Quasar 670 | LGC Biosearch Technologies | SS631302-27 |
| camkii_28: 5’- GTCTCACATAGGCAATGCAA-3’ Quasar 670 | LGC Biosearch Technologies | SS631302-28 |
| camkii_29: 5’- CATTCTGCCATTTGTTATCG-3’ Quasar 670 | LGC Biosearch Technologies | SS631302-29 |
| camkii_30: 5’- CTTATTTTGGCAGATGCACT-3’ Quasar 670 | LGC Biosearch Technologies | SS631302-30 |
| chic-570-1: 5’- CCGCAACACCGACGATTT-3’ Quasar 570 | LGC Biosearch Technologies | SS630980-01 |
| chic-570-2: 5’- CGGGGGTCACACGAAATT-3’ Quasar 570 | LGC Biosearch Technologies | SS630980-02 |
| chic-570-3: 5’- CGAGTCGCACTTTCGTTT-3’ Quasar 570 | LGC Biosearch Technologies | SS630980-03 |
| chic-570-4: 5’- TGTTGCTTTACCGCACGG-3’ Quasar 570 | LGC Biosearch Technologies | SS630980-04 |
| chic-570-5: 5’- GATCTGGATATGGATCGC-3’ Quasar 570 | LGC Biosearch Technologies | SS630980-05 |
| chic-570-6: 5’- GTCGTGGGTGCGGATTAA-3’ Quasar 570 | LGC Biosearch Technologies | SS630980-06 |
| chic-570-7: 5’- GCTCATGGTGCTTTGTTT-3’ Quasar 570 | LGC Biosearch Technologies | SS630980-07 |
| chic-570-8: 5’- GTCCACATAATCTTGCCA-3’ Quasar 570 | LGC Biosearch Technologies | SS630980-08 |
| chic-570-9: 5’- CGAGGCCAGGAGTTGGTT-3’ Quasar 570 | LGC Biosearch Technologies | SS630980-09 |
| chic-570-10: 5’- GATGCACGCCTTGGTCAC-3’ Quasar 570 | LGC Biosearch Technologies | SS630980-10 |
| chic-570-11: 5’- CCAAATGTTGCCGTCGTG-3’ Quasar 570 | LGC Biosearch Technologies | SS630980-11 |
| chic-570-12: 5’- TTTCCTCTGCTACACACA-3’ Quasar 570 | LGC Biosearch Technologies | SS630980-12 |
| chic-570-13: 5’- GCATTTTTACTCGATCCA-3’ Quasar 570 | LGC Biosearch Technologies | SS630980-13 |
| chic-570-14: 5’- CCGCTGATCAGTTTGGAG-3’ Quasar 570 | LGC Biosearch Technologies | SS630980-14 |
| chic-570-15: 5’- CCGTCCTGCTGGTCAAAG-3’ Quasar 570 | LGC Biosearch Technologies | SS630980-15 |
| chic-570-16: 5’- AGTGTCACGCCGTTGCTG-3’ Quasar 570 | LGC Biosearch Technologies | SS630980-16 |
| chic-570-17: 5’- TAAATGTACCGCTGGCCG-3’ Quasar 570 | LGC Biosearch Technologies | SS630980-17 |
| chic-570-18: 5’- CGGTCTGTGCCGGAAAGG-3’ Quasar 570 | LGC Biosearch Technologies | SS630980-18 |
| chic-570-19: 5’- GTCTTCATGCAGTGCACT-3’ Quasar 570 | LGC Biosearch Technologies | SS630980-19 |
| chic-570-20: 5’- ACGATCACGGCTTGTGTT-3’ Quasar 570 | LGC Biosearch Technologies | SS630980-20 |
| chic-570-21: 5’- GGGATCCTCGTAGATGGA-3’ Quasar 570 | LGC Biosearch Technologies | SS630980-21 |
| chic-570-22: 5’- TCTCTACCACGGAAGCGG-3’ Quasar 570 | LGC Biosearch Technologies | SS630980-22 |

| | | |
|---|---|---|
| chic-570-23: 5'- CTATTCTCCTAGTACCCG-3' Quasar 570 | LGC Biosearch Technologies | SS630980-23 |
| chic-570-24: 5'- TCATTTACGGTTCGCTCT-3' Quasar 570 | LGC Biosearch Technologies | SS630980-24 |
| chic-570-25: 5'- GTTGGTTTTTCTTTTCCC-3' Quasar 570 | LGC Biosearch Technologies | SS630980-25 |
| chic-570-26: 5'- TTTCCTCTGCTACACACA-3' Quasar 570 | LGC Biosearch Technologies | SS630980-26 |
| chic-570-27: 5'- GCATTTTTACTCGATCCA-3' Quasar 570 | LGC Biosearch Technologies | SS630980-27 |
